# Common genetic variation in humans impacts *in vitro* susceptibility to SARS-CoV-2 infection

**DOI:** 10.1101/2020.09.20.300574

**Authors:** Kristina Dobrindt, Daisy A. Hoagland, Carina Seah, Bibi Kassim, Callan P. O’Shea, Marina Iskhakova, Michael B. Fernando, P.J. Michael Deans, Samuel K. Powell, Ben Javidfar, Aleta Murphy, Cyril Peter, Rasmus Møeller, Meilin Fernandez Garcia, Masaki Kimura, Kentaro Iwasawa, John Crary, Darrell N. Kotton, Takanori Takebe, Laura M. Huckins, Benjamin R. tenOever, Schahram Akbarian, Kristen J. Brennand

## Abstract

The host response to SARS-CoV-2, the etiologic agent of the COVID-19 pandemic, demonstrates significant inter-individual variability. In addition to showing more disease in males, the elderly, and individuals with underlying co-morbidities, SARS-CoV-2 can seemingly render healthy individuals with profound clinical complications. We hypothesize that, in addition to viral load and host antibody repertoire, host genetic variants also impact vulnerability to infection. Here we apply human induced pluripotent stem cell (hiPSC)-based models and CRISPR-engineering to explore the host genetics of SARS-CoV-2. We demonstrate that a single nucleotide polymorphism (rs4702), common in the population at large, and located in the 3’UTR of the protease FURIN, impacts alveolar and neuron infection by SARS-CoV-2 *in vitro*. Thus, we provide a proof-of-principle finding that common genetic variation can impact viral infection, and thus contribute to clinical heterogeneity in SARS-CoV-2. Ongoing genetic studies will help to better identify high-risk individuals, predict clinical complications, and facilitate the discovery of drugs that might treat disease.

## INTRODUCTION

A growing number of human genetic variants have been identified that contribute to enhanced susceptibility or resistance to viral diseases (Kenney et al., 2017). Genetic discoveries related to virus–host interactions implicate genes encoding virus receptors, receptor-modifying enzymes, and a wide variety of innate and adaptive immunity-related proteins in disease outcome. Such insights have been made across a range of pathogenic viruses, including influenza A virus (Ciancanelli et al., 2015; Everitt et al., 2012), respiratory syncytial virus (Everitt et al., 2012), norovirus (Lindesmith et al., 2003), rotavirus (Payne et al., 2015), parvovirus (Hellberg et al., 2002), and human immunodeficiency virus (Samson et al., 1996).

There is marked variability between individuals in response to SARS-CoV-2 infection, with outcomes ranging from asymptomatic (∼15%), mild-moderate (50%), severe (13.8%), and critical (6.1%) (Guan et al., 2020). There is an urgent need to explore the molecular mechanisms underlying this unexpected clinical heterogeneity. For viral infections, a range of possible explanations can impact outcome: genetic predisposition (e.g. blood type (Ellinghaus et al., 2020; Zhao et al., 2020b; Zietz and Tatonetti, 2020), HLA genotype (Dutta et al., 2018), ethnicity (Das and Ghate, 2020)), immune repertoire (e.g. cross-reacting antibodies from other coronaviruses (Yuan et al., 2020)) and/or viral load (Chu et al., 2004) (e.g. reports of poor outcomes in infected medical workers). Clinical outcome is not simply a result of the adaptive immune response, as evidence is mounting that seroconversion can occur prior to symptom recovery (Wölfel et al., 2020) as well as in asymptomatic cases (Okba et al., 2020). Although the host response to SARS-CoV-2 has been defined in lung cultures and tissue (Karczewski et al., 2020), it remains unclear the full extent to which this varies across cell types and donors. The expression of a number of human genes likely modulates infection; given the complicated host immune response to SARS-CoV-2 (Blanco-Melo et al., 2020; Qin et al., 2020) and increased disease severity in patients with mutations in immune signaling receptors (van der Made et al., 2020)(Cassanova *Science*, in press), this likely includes genes involved in the innate immune system.

In an effort to determine whether host variants that may influence viral entry might contribute to the heterogeneity of COVID-19 symptoms, we assessed human variants of FURIN. The spike glycoprotein that resides on the surface of the SARS-CoV-2 virion facilitates viral entry into target cells by engaging host angiotensin-converting enzyme 2 (ACE2) as the entry receptor, and host cellular serine protease TMPRSS2 for spike protein priming (Hoffmann et al., 2020; Yan et al., 2020). More controversial is the extent to which host CD147, a transmembrane glycoprotein, serves as a secondary entry receptor (Shilts and Wright, 2020; Wang et al., 2020). The SARS-CoV-2 spike protein incorporates a four amino acid insertion that introduces a proposed cleavage site for FURIN, a host membrane-bound proprotease convertase, potentially resulting in priming of spike protein before viral exit from the cell (Coutard et al., 2020; Wrapp et al., 2020). In contrast to SARS-CoV-2, the SARS-CoV spike protein lacks this FURIN-cleavage site, thus requiring cleavage to facilitate subsequent cell entry (Belouzard et al., 2009; Walls et al., 2020). This theoretical hijacking of host FURIN activity is one possible explanation for the increased infectivity of SARS-CoV-2. Not only do these host genes show tissue-specific (Papatheodorou et al., 2020) and cell-type-specific (Franzen et al., 2019) expression patterns, but each is associated with non-coding common genetic variants thought to regulate expression of each gene (GTEx Consortium et al., 2017). Here we test the hypothesis that variability in the expression of host genes between cell types and individuals predicts susceptibility to infection. Given that CRISPR (clusters of regularly interspaced short palindromic repeats)-based allelic conversion of rs4702 in human induced pluripotent stem cells (hiPSCs) from AA to GG decreased *FURIN* mRNA levels, reduced neurite outgrowth, and altered neuronal activity in induced glutamatergic neurons (Schrode et al., 2019), here we consider the impact of *FURIN* expression and genotype on SARS-CoV-2 infection across hiPSC-derived lung, intestinal and brain models. Our findings suggest that uncovering the genetic underpinnings of SARS-CoV-2 outcomes may help predict susceptibility for COVID-19, as well as facilitate precision treatment and prevention approaches.

## RESULTS

### Infection of neurons by SARS-CoV-2 in vivo and in vitro

A subset of COVID-19 patients present with neurological symptoms (Mao et al., 2020), including headaches, dizziness, and defects in smell and taste reported in 12-60% of patients (Chen et al., 2020). COVID-19 has been associated with acute disseminated encephalomyelitis (ADEM) (Pilotto et al., 2020; Zhang et al., 2020), mid- and long-term sequels including classical Guillain–Barré syndrome (Toscano et al., 2020; Zhao et al., 2020a), and COVID-19-associated delirium (McLoughlin et al., 2020). After hospital discharge, an alarmingly high fraction of patients, as high as 33%, suffer from a dysexecutive syndrome consisting of inattention, disorientation, or poorly organized movements in response to command (Helms et al., 2020). It is critical to resolve whether potential SARS-CoV-2 pathological mechanisms include direct infection of brain cells, consistent with reports of neurotrophism and trans-synaptic spread by other coronaviruses (Desforges et al., 2014; Li et al., 2020), or simply reflect endothelial injury, vascular coagulopathy, and/or diffuse neuroinflammatory processes.

A variety of evidence reveals that human brain tissue and cultured neurons express critical host genes required for viral entry. RNA transcripts and protein for *ACE2, FURIN*, and *CD147* were detected in post-mortem adult human brain (prefrontal cortex (PFC) and midbrain substantia nigra (SN)) (**Fig. 1A,B; Supplemental Fig. 1**). Likewise, RNA-sequencing data from hiPSC-derived neural progenitor cells (NPCs) (Hoffman et al., 2017), *NGN2*-induced glutamatergic neurons (Ho et al., 2016; Zhang et al., 2013), *ASCL1/DLX2*-induced GABAergic neurons (Barretto et al., 2020; Yang et al., 2017), *ASCL1/NURR1/LMX1A*-induced dopaminergic neurons (Rehbach et al., 2020), as well as differentiated forebrain neurons (FB, comprised of a mixture of neurons and astrocytes), confirmed expression of many candidate host genes (Gordon et al., 2020; Ou et al., 2020; Wruck and Adjaye, 2020) (**Fig. 1A**). Expression of *FURIN, ACE2, CD147*, and *TMPRSS2* in *NGN2*-induced glutamatergic neurons, in mock (uninfected) and SARS-CoV-2 infected populations, was confirmed by qPCR, normalized to lung alveolosphere samples to facilitate inter-cell type comparisons of expression, with *FURIN* and *CD147* being most prominent (**Fig. 1C**).

**Figure 1.**
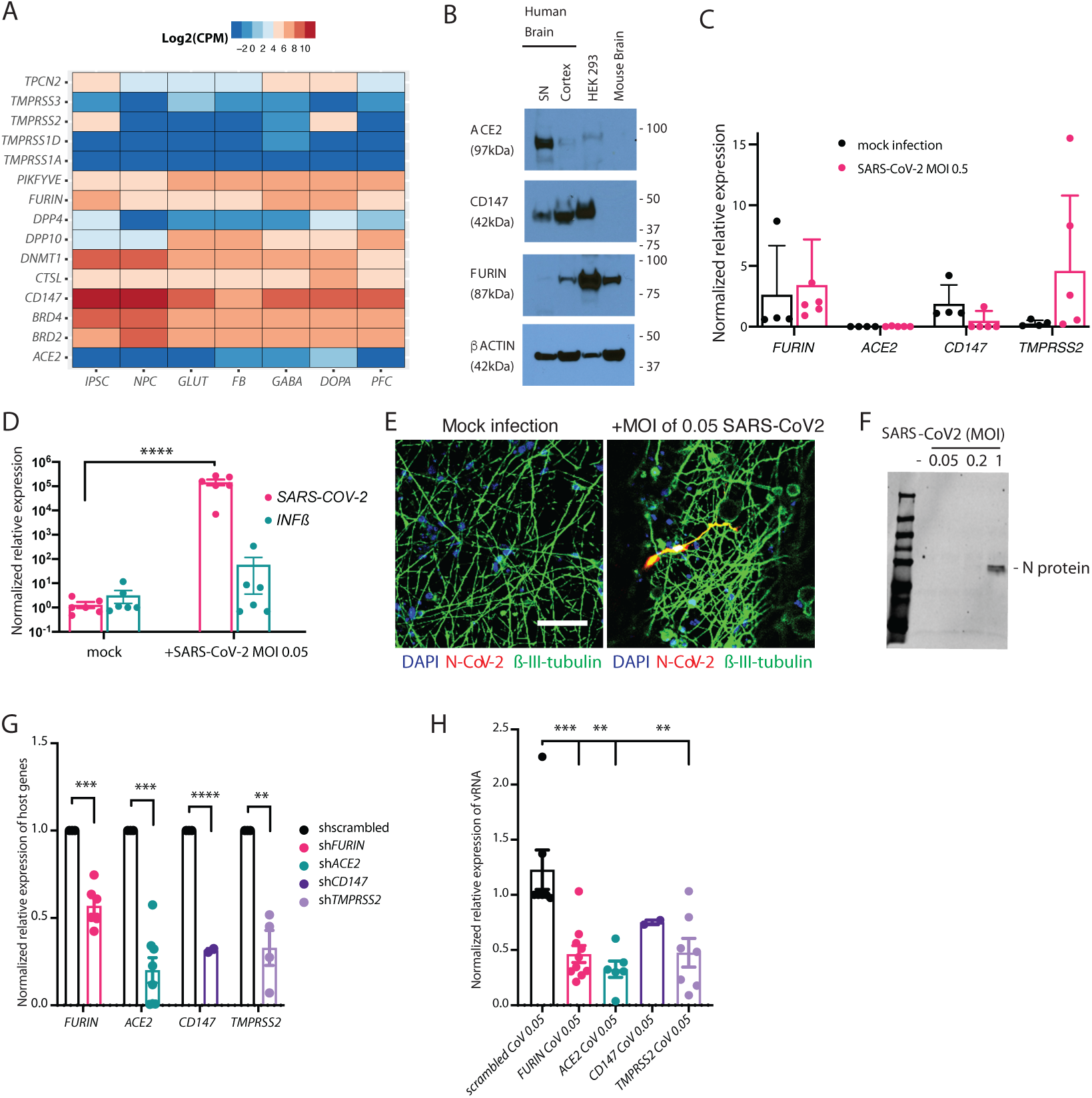
Infection of neurons by SARS-CoV-2 *in vitro*. **A**. Expression of host genes associated with SARS-CoV-2 infection (Gordon et al., 2020; Ou et al., 2020; Wruck and Adjaye, 2020) examined by RNA-sequencing (Log 2 RNA-seq, CPM medians) of post-mortem human prefrontal cortex (PFC), human induced pluripotent stem cells (hiPSCs), neural progenitor cells (NPCs) and four hiPSC-derived neuronal cell types: induced glutamatergic (GLUT), differentiated forebrain (FB) populations (comprised of glutamatergic neurons, GABAergic neurons and astrocytes), induced GABAergic (GABA) and induced dopaminergic (DOPA). **B**. Immunoblot showing ACE2, FURIN and CD147 protein in post-mortem human adult substantia nigra (SN) and PFC. **C**. Baseline host gene expression in d21 *NGN2*-induced glutamatergic neurons, uninfected or infected with SARS-CoV-2 (MOI 0.5), normalized relative to alveolospheres, to ensure comparability across cell types. **D**. qPCR analysis of relative change in subgenomic vRNA N transcript and Interferon β (IFNβ) in mock and SARS-CoV-2 infected d21 *NGN2*-induced glutamatergic neurons (MOI of 0.05, 24 hours). **E**. Representative immunostaining for SARS-CoV-2 nucleocapsid (N) protein (red), neuronal marker β-III-tubulin (green) and DAPI (blue) in d7 *NGN2*-induced glutamatergic neurons infected with a SARS-CoV-2 (MOI of 0.05, 48 hours). Scale bar: 50μm **F**. Immunoblot against SARS-CoV-2 nucleocapsid (N) protein shows replication in *NGN2*-induced glutamatergic neurons at a MOI of 1 after 24 hours. **G**. shRNA knock down of *FURIN, ACE2, CD147*, and *TMPRSS2* host genes, normalized to 18S levels and scrambled, in d7 *NGN2*-induced glutamatergic neurons. T-test, **p < 0.01, ***p < 0.001, ****p < 0.0001. **H**. Subgenomic vRNA nucleocapsid transcript in infected *NGN2*-induced glutamatergic neurons (MOI of 0.05 for 24 hours) treated with shRNAs against SARS-CoV-2 receptors. T-test, **p < 0.01, ***p < 0.001.

To determine neuronal susceptibility to SARS-CoV-2 virus *in vitro*, human *NGN2*-induced glutamatergic neurons derived from two independent donors were infected with SARS-CoV-2 virus at low multiplicity of infection (MOI 0.05). Glutamatergic neurons were susceptible to SARS-CoV-2 virus infection, rapidly increasing expression of subgenomic viral RNA (vRNA) nucleocapsid transcript by qPCR (148,124-fold, p-value <0.0001) within 24 hours (**Fig. 1D**) and protein immunohistochemistry for SARS-CoV-2 nucleocapsid (N) within 48 hours of infection (MOI of 0.05) (**Fig. 1E**). At higher titers (MOI 1, 24 hours), immunoblot against N protein was consistent with SARS-CoV-2 replication in neurons (**Fig. 1F**).

To further explore the susceptibility of the *in vivo* brain to SARS-CoV-2, we conducted RNA-sequencing of brains of 3-5 week old male Golden Syrian hamsters infected intranasally with SARS-CoV-2 or a phosphate buffered saline (PBS) control (Imai et al., 2020; Si et al., 2020). By RNA sequencing, we identified high levels of SARS-CoV-2 host gene expression in hamster brain tissue, with notably high levels of *FURIN* and *BSG/CD147* (**Fig. 2A)**. SARS-CoV-2 infection resulted in widespread up- and downregulation of transcripts (**Fig. 2B**), notably upregulation of interferon-associated genes within 24 hours, an effect that subsided somewhat by day 8 (**Fig. 2C)**. Gene set enrichment analysis of differentially expressed genes was performed across a collection of 698 neural-themed gene sets subdivided into 8 categories (Schrode et al., 2019). Notably, significantly upregulated gene sets (false discovery rate (FDR) < 5%) included those related to abnormal neuronal morphology and development, membrane trafficking and abnormal synaptic transmission (clustered hierarchically by significance in (**Fig. 2D)**, while downregulated gene sets included ion channel and neurotransmitter signaling pathways (**Fig. 2E**). Altogether, these analyses are consistent not just with neuronal infection by SARS-CoV-2, but further suggest that impaired neural function may occur as a result.

**Figure 2.**
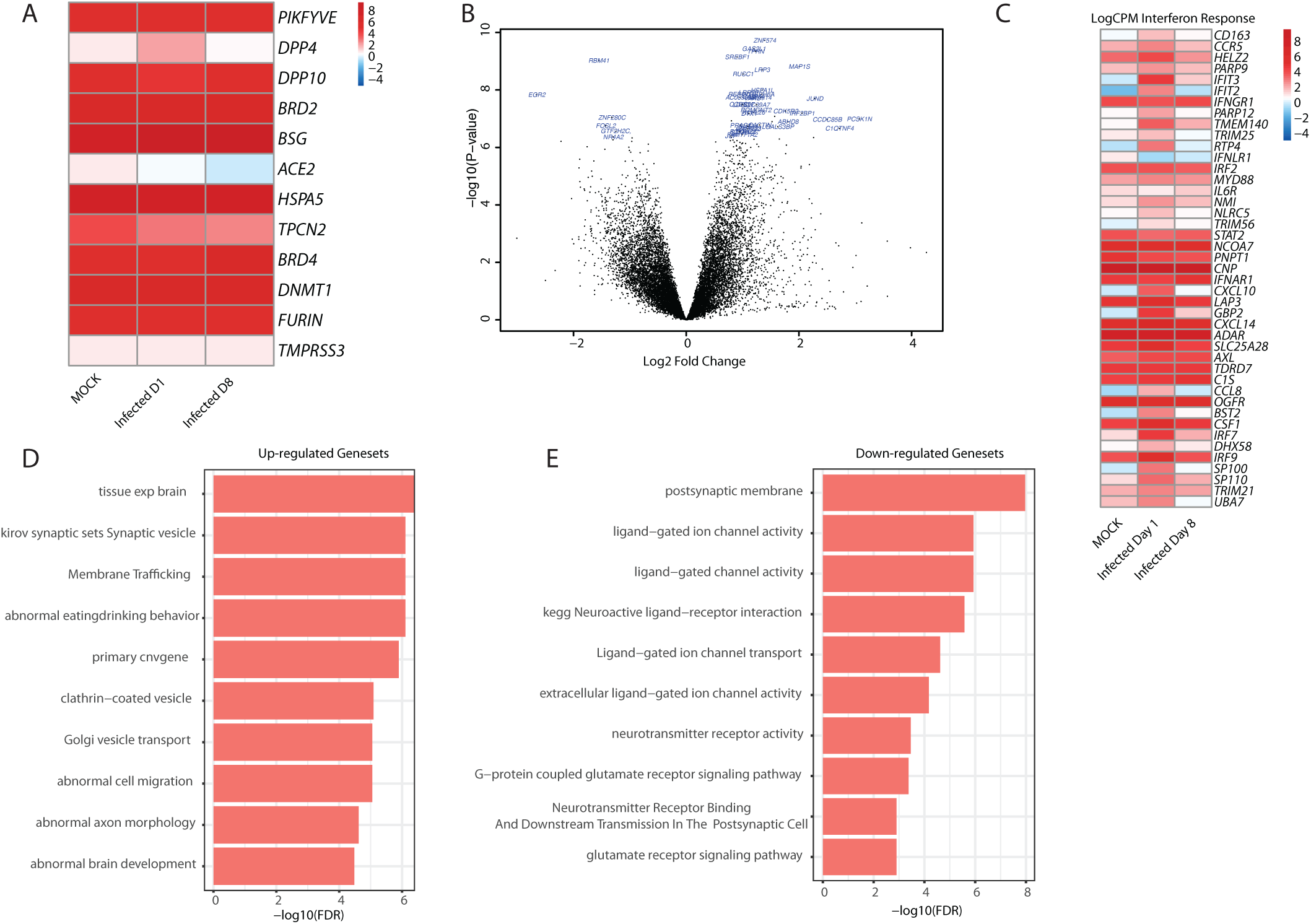
RNA-sequencing analysis of expression changes in hamster brain after SARS-CoV-2 infection *in vivo*. **A**. Expression of relevant host genes associated with SARS-CoV-2 infection in hamster brain after mock infection, 1 or 8 days following SARS-CoV-2 infection. **B**. Volcano plot highlighting the top 50 significantly differentially expressed genes after a high pfu challenge with SARS-CoV-2. **C**. Interferon response over course of infection, expression shown in log2CPM, gene set from (Blanco-Melo et al., 2020). **D**. Gene set enrichment for upregulated pathways using the MAGMA gene set. **E**. Gene set enrichment for downregulated pathways using curated gene sets (Schrode et al., 2019).

### Host gene-dependent infection of brain, lung and intestinal cells by SARS-CoV-2

SARS-CoV-2 infects hiPSC-derived lung alveolospheres (Abo et al., 2020; Han et al., 2020; Huang et al., 2020) and intestinal organoids (Lamers et al., 2020; Yang et al., 2020; Zhou et al., 2020), particularly after dissociation; here we similarly apply published step-wise differentiation protocols for distal lung alveolar type 2-like epithelium (hereafter “alveolar cells”) (Jacob et al., 2019) and intestinal organoids (Koike et al., 2019; Zhang et al., 2018).

Alveolospheres formed after 30 days (**Fig. 3A**) express host genes associated with SARS-CoV-2 infection (**Fig. 3B**). Consistent with other studies, we observed infection by SARS-CoV-2 of EPCAM and NKX2.1-positive lung alveolospheres derived from two donors across three infections on protein level (MOI 0.5, 24 hours) (**Fig. 3C)** and on RNA level (MOI 0.1 and 0.5, 24 hours) **(Fig.3D**). Dissociated alveolar cells showed detectable SARS-CoV-2 N protein (MOI of 0.1, 24 hours) (**Fig. 3E**). Similar to alveolospheres, d50 EPCAM-positive intestinal organoids (one donor) also showed SARS-CoV-2 infection on protein level (MOI 0.5 and 1) (**Fig. 4A)** and on RNA level (MOI 0.05 and 0.5, 24 hours) (Fig. 4**C**). Intestinal organoids express high levels of *FURIN, ACE2* and *CD147*, but lower levels of *TMPRSS2*, relative to alveolospheres (**Fig. 4B**). We optimized MOIs (neuron: 0.05, alveolosphere: 0.1, intestine: 0.05) and exposure times (neuron: 24 hours for RNA, 48 hours for protein; alveolosphere: 24 hours; intestine: 24 hours for RNA, 48 hours for protein) for subsequent studies of host gene and variant effects, towards maximizing cell viability and detection of SARS-CoV-2 nucleocapsid (N) protein.

**Figure 3.**
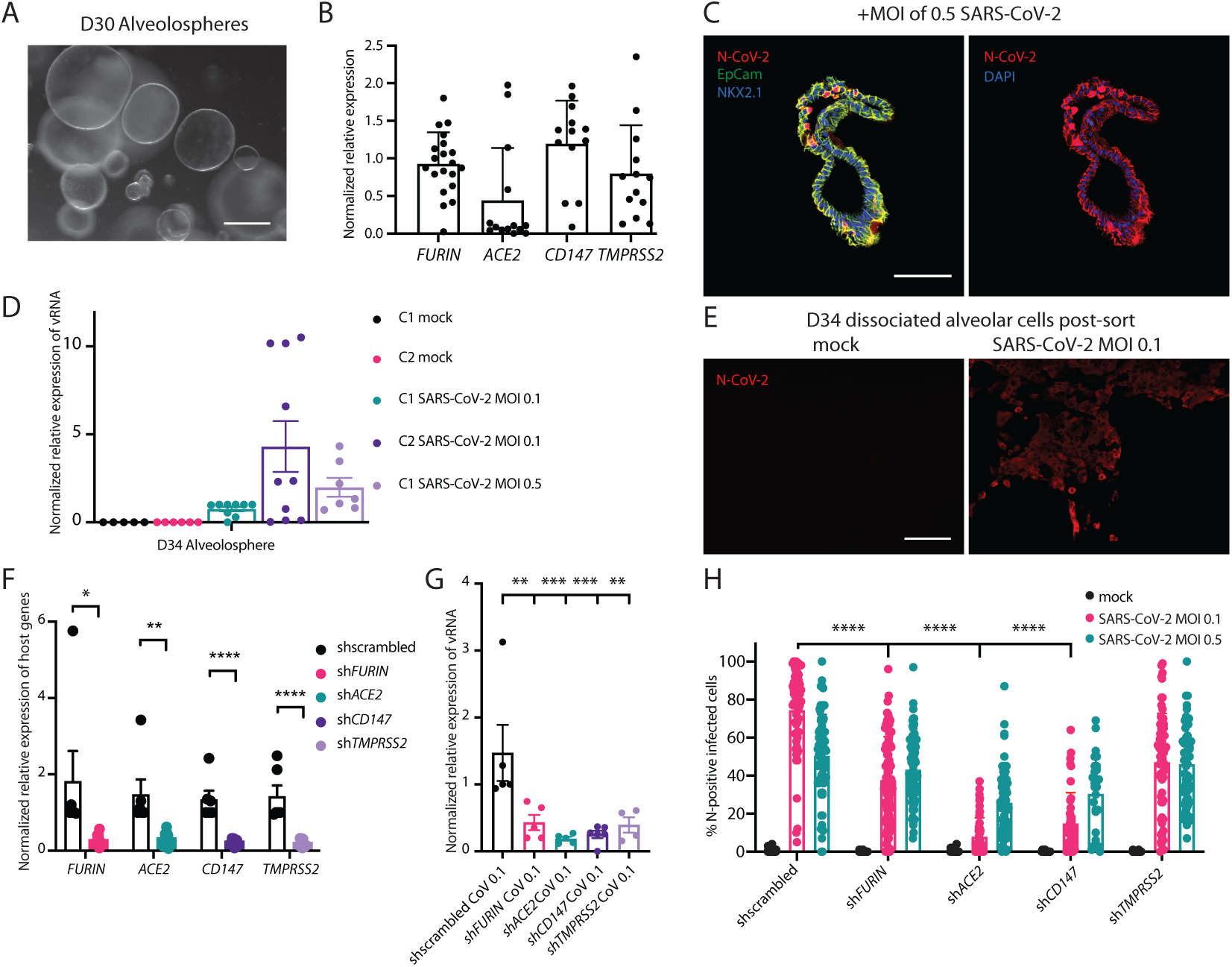
Infection of lung alveolospheres by SARS-CoV-2 *in vitro*. **A**. Representative brightfield image of alveolosphere cultures on d30. 100μm. **B**. Baseline SARS-CoV-2 host gene expression in alveolospheres. Scale bar: 100μm. **C**. Representative immunofluorescence staining against SARS-CoV-2 nucleocapsid (N) (red), epithelial marker EPCAM (green), Nkx2.1 (blue, left) and DAPI (blue, right) in alveolospheres with mock or SARS-CoV-2 infection (MOI 0.5, 24 hours). Scale bar: 100μm. **D**. qPCR analysis of relative change in subgenomic vRNA N transcript in mock and SARS-CoV-2 infected d34 alveolospheres from two donors (MOI of 0.1 or 0.5, 24 hours). **E**. Dissociated alveolar cells after SARS-CoV-2 infection on day 34 are positive for SARS-CoV-2 N protein (red). Scale bar: 100μm. **F**. shRNA knock down of *FURIN, ACE2, CD147*, and *TMPRSS2* host genes, normalized to 18S levels and C1 alveolosphere (MOI 0.1, 24 hours). T-test, **p < 0.01, ****p < 0.0001, error bars show SEM. **G**. qPCR analysis of relative change in subgenomic vRNA N transcript in mock and SARS-CoV-2 infected dissociated alveolar cells expressing shRNAs against *ACE2, TMPRSS2, CD147* and *FURIN*, as well as a scrambled control, **p < 0.01, ***p < 0.001, error bars shown as SEM. **H**. Quantification of high-content imaging of for SARS-CoV-2 nucleocapsid (N) protein in mock and SARS-CoV-2 infected dissociated alveolar cells expressing shRNAs against *ACE2, TMPRSS2, CD147* and *FURIN*, as well as a scrambled control. SARS-CoV-2 infected at MOI of 0.1 and 0.5 for 24 hours. Each data point represents at least 100 cells. 2-way ANOVA; ****p < 0.0001.

**Figure 4.**
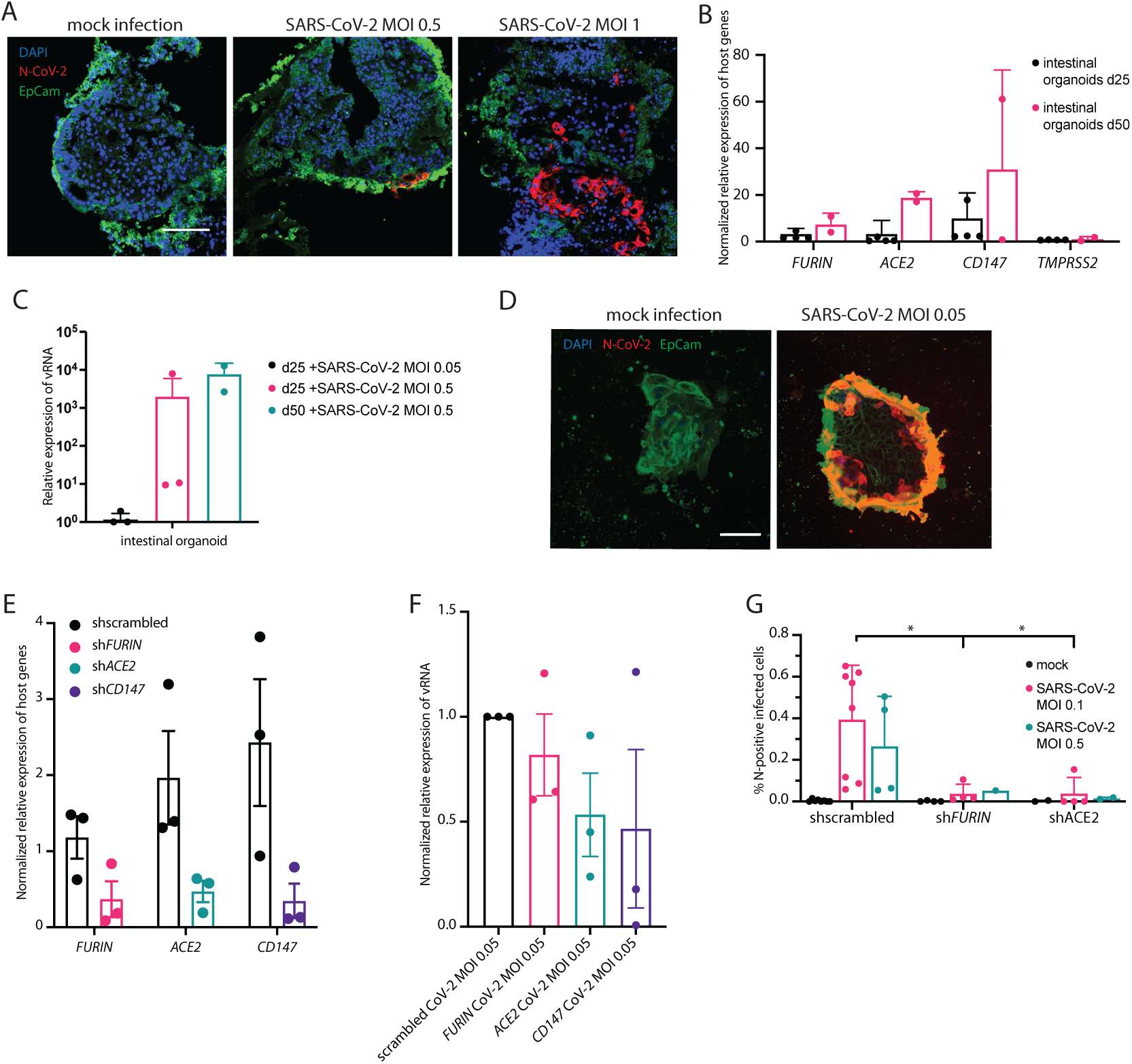
Infection of intestinal organoids by SARS-CoV-2 *in vitro*. **A**. Baseline SARS-CoV-2 host gene expression in mock treated **A**. Representative immunofluorescence staining of intestinal organoids for SARS-CoV-2 nucleocapsid (N) (red), the epithelial marker EPCAM (green) and DAPI (blue), following mock or SARS-CoV-2 infection (MOI of 0.5 or 1, 48 hours). Scale bar: 100μm. **B**.intestinal organoids day 24 and d50. **C**. qPCR analysis of relative change in subgenomic vRNA N transcript in mock and SARS-CoV-2 infected intestinal organoids infected (MOI of 0.05 or 0.5, 24 hours). **D**. Representative immunofluorescence staining of dissociated intestinal cells after SARS-CoV-2 infection on day 25 (MOI of 0.05, 48 hours), positive for SARS-CoV-2 N protein (red), EPCAM (green), DAPI (blue). Scale bar: 50μm. **E**. shRNA knock down of *FURIN, ACE2*, and *CD147* host genes in dissociated intestinal cells, normalized to 18S levels and infected scrambled control. **F**. qPCR analysis of relative change in subgenomic vRNA N transcript in mock and SARS-CoV-2 infected dissociated intestinal cells expressing shRNAs against *ACE2, CD147* and *FURIN*, as well as a scrambled control, error bars shown as SEM. **G**. Quantification of high-content imaging for SARS-CoV-2 nucleocapsid (N) protein in mock and SARS-CoV-2 infected dissociated intestinal cells expressing shRNAs against *ACE2* and *FURIN*, as well as a scrambled control. (SARS-CoV-2 infected at MOI of 0.05 and 0.5 for 48 hours.) Each data point represents one 96-well. 2-way ANOVA; *p < 0.05.

Alveolospheres formed after 30 days (**Fig. 3A**) express host genes associated with SARS-CoV-2 infection (**Fig. 3B**). Consistent with other studies, we observed infection by SARS-CoV-2 of EPCAM and NKX2.1-positive lung alveolospheres derived from two donors across three infections on protein level (MOI 0.5, 24 hours) (**Fig. 3C)** and on RNA level (MOI 0.1 and 0.5, 24 hours) **(Fig.3D**). Dissociated alveolar cells showed detectable SARS-CoV-2 N protein (MOI of 0.1, 24 hours) (**Fig. 3E**). Similar to alveolospheres, d50 EPCAM-positive intestinal organoids (one donor) also showed SARS-CoV-2 infection on protein level (MOI 0.5 and 1, 48 hours) (**Fig. 4A)** and on RNA level (MOI 0.05 and 0.5, 24 hours) (Fig. 4**C**). Intestinal organoids express high levels of *FURIN, ACE2* and *CD147*, but lower levels of *TMPRSS2*, relative to alveolospheres (**Fig. 4B**). Dissociated intestinal cells showed detectable SARS-CoV-2 N protein (MOI of 0.05, 48 hours) (**Fig. 4D**). We optimized MOIs (neuron: 0.05, alveolosphere: 0.1, intestine: 0.05) and exposure times (neuron: 24 hours for RNA, 48 hours for protein; alveolosphere: 24 hours; intestine: 24 hours for RNA, 48 hours for protein) for subsequent studies of host gene and variant effects, towards maximizing cell viability and detection of SARS-CoV-2 nucleocapsid (N) protein.

To test the functional impact of decreasing expression of critical host genes, we applied a shRNA strategy to knock down expression of *ACE2, FURIN, CD147* and *TMPRSS2* in neurons (**Fig. 1G:** *FURIN* (expression reduced to 56.9%, p-value <0.00045), *ACE2* (expression reduced to 20%, p-value 0.0001), *CD147* (expression reduced to 31.5%, p-value <0.0001) and *TMPRSS2* (expression reduced to 32.9%, p-value 0.002374)), lung alveolar cells (**Fig. 3F:** *FURIN* (expression reduced to 14.5%, p-value 0.0208), *ACE2* (expression reduced to 21.1%, p-value 0.002), *CD147* (expression reduced to 16.5%, p-value <0.0001) and *TMPRSS2* (expression reduced to 13.8%, p-value <0.0001)) and intestinal (**Fig. 4E:** *ACE2* (expression reduced to 23.9%, p-value 0.0768), *FURIN* (expression reduced to 31.3%, p-value 0.0898), and *CD147* (expression reduced to 14.2%, p-value 0.0732)). Observed knockdown did not significantly differ across cell types or donors. Moreover, susceptibility to infection was quantified by qPCR for subgenomic vRNA nucleocapsid transcript and immunocytochemistry with high-content imaging for N protein. By comparing vRNA nucleocapsid transcript expression in mock and SARS-CoV-2 infected neurons (**Fig. 1H**), dissociated alveolosphere cells (**Fig. 3G**), and dissociated intestinal organoid cells (**Fig. 4F)** expressing shRNAs against *ACE2, TMPRSS2, CD147* and *FURIN*, as well as a scrambled control, we demonstrate non-cell-type-specific requirements for *ACE2, CD147* and *FURIN:* (**Fig. 1H:** *FURIN* (vRNA expression reduced to 37.7%, p-value 0.0005), *ACE2* (vRNA expression reduced to 26.6%, p-value 0.0011), *CD147* (vRNA expression reduced to 61.1%, p-value 0.2162) and *TMPRSS2* (vRNA expression reduced to 38.7%, p-value 0.0053)), lung alveolar cells (**Fig. 3G:** *FURIN* (vRNA expression reduced to 29.3%, p-value 0.0055), *ACE2* (vRNA expression reduced to 12.8%, p-value 0.0008), *CD147* (vRNA expression reduced to 17.2%, p-value 0.0009) and *TMPRSS2* (vRNA expression reduced to 26.8%, p-value 0.0067)) and intestinal (**Fig. 4F:** *ACE2* (vRNA expression reduced to 53.3%, p-value 0.0784), *FURIN* (vRNA expression reduced to 81.9%, p-value 0.4035), and *CD147* (vRNA expression reduced to 46.6%, p-value 0.2297)). These findings were confirmed through quantification of fluorescence via high-content imaging in dissociated alveolar cells for N protein, which revealed decreased total infected cells compared to scrambled control (MOI of 0.1, 24 hours): sh*FURIN* (N protein reduced to 50.7%, p-value <0.0001), sh*ACE2* (N protein reduced to 10.7%, p-value <0.0001), sh*CD147* (N protein reduced to 20.1%, p-value <0.0001) and *TMPRSS2* (N protein reduced to 63.2%, p-value <0.0001) (**Fig. 3H**). Consistent with this, knockdown of *FURIN* and *ACE2* in dissociated organoid intestinal cells led to a decreased number of N-positive SARS-CoV-2 infected cells through quantification of fluorescence via high-content imaging (MOI 0.05 after 48 hours): sh*FURIN* (N protein reduced to 9.5%, p-value 0.0197), and sh*ACE2* (N protein reduced to 9.8%, p-value 0.0203) (**Fig. 4G**).

Critically, shRNA-mediated knockdown of host genes resulted in large changes in their endogenous expression, which may not well model subtle changes in gene expression resulting from common variation between individual people but can serve here as positive controls to negatively affect SARS-CoV-2 entry and egress.

### Host FURIN rs4702 genotype-infection by SARS-CoV-2

To achieve more physiologically relevant changes in gene expression, CRISPR-based allelic conversion was used to generate isogenic hiPSCs that differed at a single non-coding regulatory single nucleotide polymorphism (SNP) at the *FURIN* locus (rs4702, NC_000015.10:g.90883330G>A). This SNP was predicted to be a quantitative trait loci (eQTL) in the human brain (Fromer et al., 2016) and *NGN2*-induced glutamatergic neurons (Forrest et al., 2017), and fine-mapping analysis identified a single putative causal cis-eQTL (probability = 0.94) (Schrode et al., 2019). rs4702 was empirically validated to regulate *FURIN* expression, neuronal outgrowth and neuronal activity following allelic conversion from AA to GG in *NGN2*-induced glutamatergic neurons (Schrode et al., 2019). Here we evaluate the regulatory effect of this common SNP, determining the extent to which rs4702 regulates *FURIN* expression and SARS-CoV-2 infection in lung cells and neurons (**Fig. 5**).

**Figure 5.**
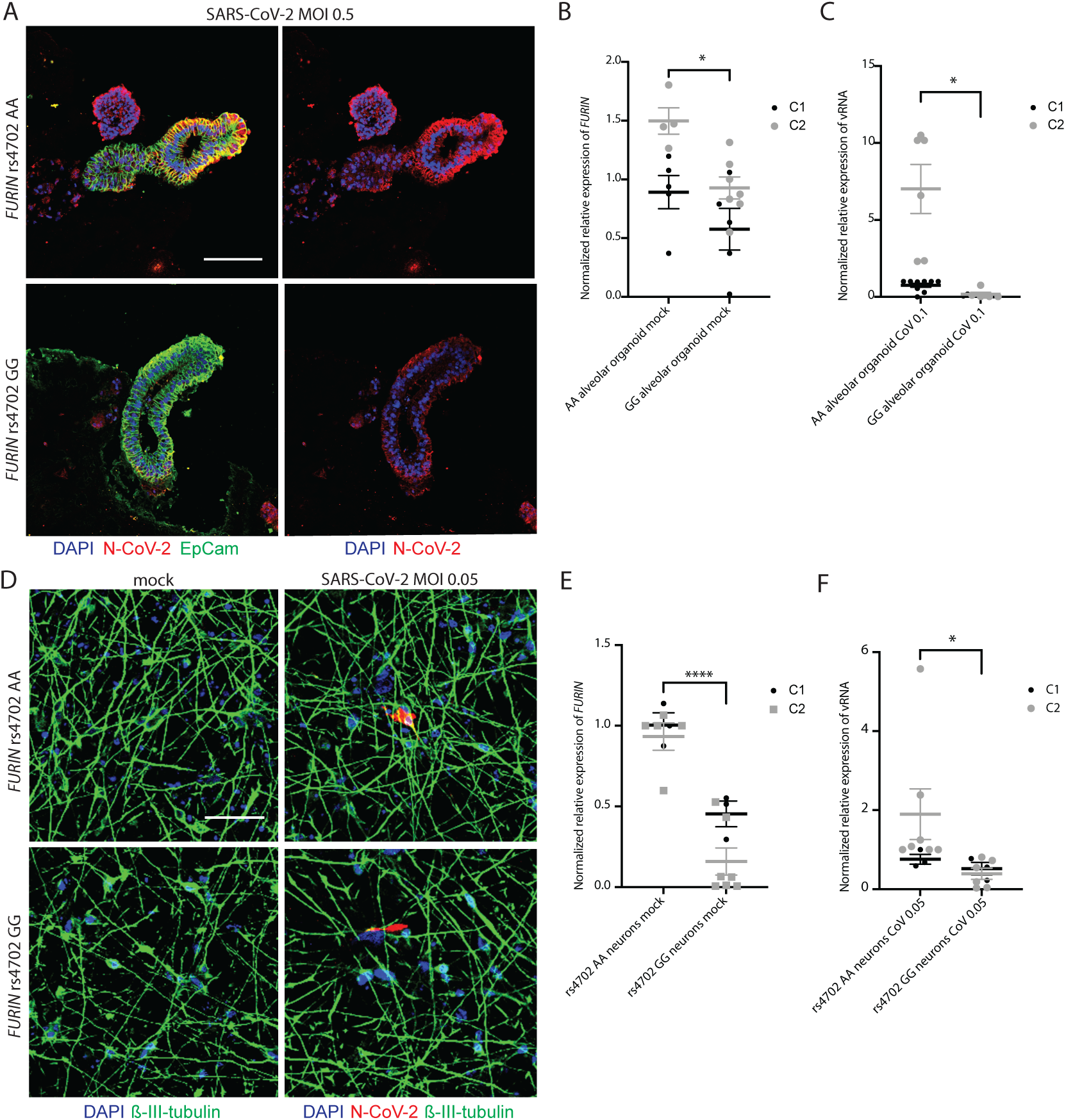
Allelic conversion at *FURIN* rs4702 in alveolospheres and neurons impacts SARS-CoV-2 infection. **A**. Representative immunofluorescence staining against SARS-CoV-2 nucleocapsid (N) protein (red), epithelial marker EPCAM (green), and DAPI (blue). Alveolospheres were generated from C2 *FURIN* rs4702 AA and GG lines and infected with mock or a MOI of 0.5 SARS-CoV-2 for 24 hours. Scale bar: 100 μm. **B**. *FURIN* RNA expression in rs4702 alveolospheres, relative to isogenic AA alveolospheres, in two control donors. Unpaired t-test (−0.327-fold, p-value 0.0347), *p < 0.05, error bars shown as SEM. **C**. qPCR analysis of relative change in subgenomic vRNA N transcript in mock and SARS-CoV-2 infected GG and AA alveolospheres from two control donors. Unpaired t-test (−0.95-fold, p-value 0.0215), *p < 0.05, error bars shown as SEM. **D**. Representative immunofluorescence staining of *NGN2*-induced glutamatergic *FURIN* rs4702 AA and GG neurons against nucleocapsid (N) protein (red), β-III-tubulin (green), and DAPI (blue), after infection SARS-CoV-2 (MOI of 0.05, 48 hours). Scale bar: 50μm. **E**. *FURIN* RNA expression in d7 rs4702 GG *NGN2*-induced neurons, relative to isogenic AA neurons. Unpaired t-test (−0.74-fold, p-value <0.0001) ****p < 0.0001, error bars shown as SEM. **F**. qPCR analysis of relative change in subgenomic vRNA N transcript in mock and SARS-CoV-2 infected d7 *NGN2*-induced neurons (MOI of 0.05, 24 hours). Unpaired t-test (−0.72-fold, p-value 0.0416), *p < 0.05, error bars shown as SEM.

Strikingly, rs4702 GG alveolospheres exhibit a reduced *FURIN* expression (−0.327-fold, p-value 0.0347) (**Fig. 5B**) and demonstrated reduced genotype-dependent SARS-CoV-2 infection across both donors (−0.95-fold, p-value 0.0215) (**Fig. 5A,C**). In 7-day-old CRISPR-edited rs4702 GG *NGN2*-induced glutamatergic neurons, we observed decreased *FURIN* expression relative to AA neurons, from two independent donors (−0.74-fold, p-value <0.0001) (**Fig 5E**), consistent with (Schrode et al., 2019). Both rs4702 AA and rs4702 GG neurons were infected by SARS-CoV-2 (MOI 0.05) at 48 hours (**Fig. 5D**), trending towards decreased SARS-CoV-2 infection in GG neurons by qPCR for subgenomic vRNA nucleocapsid transcript after 24 hours (−0.72-fold, p-value 0.0416) (**Fig. 5F**). Altogether, these results indicate that reduced *FURIN* levels mediated by rs4702 result in decreased SARS-CoV-2 infection.

Overall, these findings suggest not only that gross manipulations of host genes can impact SARS-CoV-2 infection in lung and brain cells, but also that the more subtle gene expression changes associated with genetic common variants are sufficient to impact viral infection levels.

## DISCUSSION

We apply a hiPSC-based model to explore the genetics of cell-type-specific host response to SARS-CoV-2. First, we report infection of post-mitotic human neurons by SARS-CoV-2. Second, we further demonstrate functional validation of the impact of host genes (*ACE2, FURIN, CD147*, and *TMPRSS2)* and genetic variation (rs4702) on SARS-CoV-2 infection in human neurons, lung and intestinal cells. Our results highlight the importance of *FURIN* as a mediator for SARS-CoV-2 infection and further demonstrate that a common variant, rs4702, located in the 3’-UTR of the *FURIN* gene, and with allele frequency estimates from G=0.34 to 0.45 (Tryka et al., 2014), is capable of influencing SARS-CoV-2 infection *in vitro*. More specifically, CRISPR/Cas9-mediated allelic conversion (from AA to GG) at the common variant rs4702 resulted in decreased neuronal and alveolar expression of the *cis*-gene target *FURIN*, and reduced SARS-CoV-2 infection. Our isogenic hiPSC-based strategy provides a proof-of-principle demonstration that common human genetic variation may directly impact SARS-CoV-2 infection.

The neuroinvasion and neurotrophism by SARS-CoV-2 observed here is consistent with other pre-print studies, which include reports of infection of human neurons and astrocytes using monolayer and organoid approaches (Jacob et al., 2020; Ramani et al., 2020; Song et al., 2020), highlighting greater and productive infection in choroid plexus epithelial cells (Jacob et al., 2020) and less infection in NPCs (Ramani et al., 2020; Song et al., 2020). Consequences of neuronal infection include transcriptional dysregulation indicative of an inflammatory response (Jacob et al., 2020), missorted and phosphorylated Tau (Ramani et al., 2020), and increased cell death (Jacob et al., 2020; Ramani et al., 2020; Song et al., 2020).

Large-scale international consortia have assembled to identify host-genetic associations with COVID-19 clinical outcomes (Initiative, 2020; Shelton et al., 2020; Tanigawa and Rivas, 2020), although cohorts remain under-powered by genome-wide association study (GWAS) standards. To date, such studies have only found a handful of loci to be genome-wide significant (Initiative, 2020; Kachuri et al., 2020; Ramlall et al., 2020). The largest data-freeze to date (https://www.covid19hg.org) does not include rs4702 within summary statistics; however, we note an excess of nominally significant genetic variants within the *FURIN* cis-region (**Supplemental Table 1**; binomial test p-value=3.57⨯10^−31^) when comparing individuals hospitalized from COVID-19 to the general population (Initiative, 2020). This indicates the necessity of expanding GWAS to more comprehensively explore the polygenic architecture for COVID-19 clinical outcome. Such discoveries could help to generate hypotheses for drug repurposing, identify individuals at unusually high or low risk, and contribute to global knowledge of the biology of SARS-CoV-2 infection and disease. Discovery-based approaches to broadly identify all genes required for host infection and survival across tissues and cell types are also necessary to inform *in silico* drug repurposing screens (Han et al., 2020). Integration of ongoing clinical GWAS with *in vitro* CRISPR-based forward genetic screening (e.g. Perturb-seq (Dixit et al., 2016) and ECCITE-seq (Mimitou et al., 2019)) and pooled eQTL screens (e.g. crisprQTL mapping (Gasperini et al., 2019) and Census-seq (Cuomo et al., 2020; Jerber et al., 2020; Mitchell et al., 2020; Neavin et al., 2020)) approaches will more rapidly identify host variants that impact the entry, replication and egress of SARS-CoV-2, cellular survival, and immune response.

This study focused only on host genes known to mediate SARS-CoV-2 infection. The broad requirement of *ACE2* and *TMPRSS2* expression for SARS-CoV-2 infection across cell types was consistent with established literature (Hoffmann et al., 2020), whereas the observed impact of *CD147* expression was more surprising given recent evidence suggesting that CD147 is incapable of binding SARS-CoV-2 (Shilts and Wright, 2020). Together with the data shown here, this suggests that CD147 protein may enhance viral entry by means other than directly binding to or modifying SARS-CoV-2, that it may require a secondary cofactor, and/or that CD147 has cell-type-specific functions *in vivo*. The impact of *FURIN* expression was not unexpected (Coutard et al., 2020; Wrapp et al., 2020); however, because rs4702 lies in the binding site for microRNA-338 (Hou et al., 2018), this is a context-dependent eQTL, and so dependent upon cell-type-specific and donor-dependent expression of miR-338 (Schrode et al., 2019). Overall, it remains possible that our *in vitro* findings will not predict clinical response because of the influence of other genetic and/or environmental factors. For example, SARS-CoV-2 antagonism of the immune response has been substantially defined (Konno et al., 2020; Lei et al., 2020; Thoms et al., 2020), and so genetic variants that result in decreased replication efficiency of SARS-CoV-2 *in vitro* may ultimately prove advantageous for the virus *in vivo*, permitting low levels of virus to spread successfully while remaining undetected by the immune system.

The expected clinical response to SARS-CoV-2 infection in the donors from whom we derived hiPSCs is not known, therefore we cannot explore the extent to which *in vitro* findings recapitulate clinical outcomes. Moreover, hiPSC-derived cells of most (if not all) lineages are immature relative to those in adult humans (Hoffman et al., 2017; Jacob et al., 2019; Pavlovic et al., 2018), which may impact SARS-CoV-2 infection/response, particularly since clinical outcomes in children are generally mild (Ludvigsson, 2020). Given that clinical evidence suggests that SARS-CoV-2 infection can lead to heterogeneous acute and chronic outcomes across a range of tissues, future work should explore across the wider range of cell types, both the cell autonomous response to infection as well as non-cell-autonomous effects resulting from subsequent cytokine storms that occur as part of response to infection. Moving forward, we should strive towards an expanded and unbiased isogenic strategy, that will prove more broadly useful in exploring the impact of host genome on SARS-CoV-2 infection across diverse cell types and tissues.

In summary, we demonstrate that a single non-coding SNP is sufficient to impact SARS-CoV-2 infection in human neurons and alveolar cells. This work supports ongoing efforts to discover host genes associated with SARS-CoV-2 infection, both *in vitro* and in the clinic. Our hope is that such efforts might better predict clinical outcomes before the onset of symptoms and facilitate the discovery of drugs that might prevent or treat COVID-19 disease.

## MATERIALS AND METHODS

### hiPSC lines and culture

Human induced pluripotent stem cells (hiPSCs) were cultured in StemFlex media (Gibco, #A3349401) and passaged with EDTA (Life Technologies #15575-020). All wildtype and shRNA experiments in lung alveolosphere cells, as well as all CRISPR-engineered AA and GG experiments were conducted in NSB3113 (female, 18 years old, European descent) and NSB3188 (female, 14 years old, European descent) (Schrode et al., 2019). shRNA experiments in neurons were conducted in control NSB553 (male, 31 years old, European descent) and NSB2607 (male, 15 years old, European descent) hiPSCs (Hoffman et al., 2017); wildtype and shRNA intestinal cell experiments were conducted in 1383D6 (male, 36 years, Asian descent) (Takayama et al., 2017).

### hiPSC-NPC culture

hiPSC-NPCs were cultured in hNPC media (DMEM/F12 (Life Technologies #10565), 1x N-2 (Life Technologies #17502-048), 1x B-27-RA (Life Technologies #12587-010), 20 ng/mL FGF2 (Life Technologies)) on Matrigel (Corning, #354230). hiPSC-NPCs at full confluence (1-1.5⨯10^7^ cells / well of a 6-well plate) were dissociated with Accutase (Innovative Cell Technologies) for 5 mins, spun down (5 mins X 1000g), resuspended and seeded onto Matrigel-coated plates at 3-5⨯10^6^ cells / well. Media was replaced every 24 days for 4 to 7 days until next split.

### NGN2-glutamatergic neuron induction (Ho et al., 2016; Zhang et al., 2013)

#### i) Neurons d21 (Fig. 1B,C)

hiPSCs were dissociated with Accutase Cell Detachment Solution (Innovative Cell Technologies, # AT-104), counted and transduced with rtTA (Addgene 20342) and *NGN2* (Addgene 99378) lentiviruses in StemFlex media containing 10 μM Thiazovivin (Millipore, #S1459). They were subsequently seeded at 1⨯10^6^ cells/well in the prepared 6-well plate. On day 1, medium was switched to non-viral induction medium (DMEM/F12 (Thermofisher, #10565018), 1% N-2 (Thermofisher, #17502048), 2% B-27-RA (Thermofisher, #12587010)) and doxycycline (dox) was added to each well at a final concentration of 1 μg/mL. At day 2, transduced hiPSCs were treated with 500 μg/mL G418 (Thermofisher, #10131035). At day 4, medium was replaced including 1 μg/mL dox and 4 μM cytosine arabinoside (Ara-C) to reduce the proliferation of non-neuronal cells. On day 5, young neurons were dissociated with Accutase Cell Detachment Solution (Innovative Cell Technologies, # AT-104), counted and seeded at a density of 1⨯10^6^ per well of a Matrigel-coated 12-well plate. Medium was switched to Brainphys neuron medium (Brainphys (STEMCELL, # 05790), 1% N-2, 2% B--27-RA, 1 μg/mL Natural Mouse Laminin (Thermofisher, # 23017015), 10 ng/mL BDNF (R&D, #248), 10 ng/mL GDNF (R&D, #212), 500 μg/mL Dibutyryl cyclic-AMP (Sigma, #D0627), 200 nM L-ascorbic acid (Sigma, # A4403)). For seeding, 10 μM Thiazovivin (Millipore, #S1459), 500 μg/mL G418 and 4 μM Ara-C and 1 μg/mLdox were added. At day 6, medium was replaced with Brainphys neuron medium with 4 μM Ara-C and 1 μg/mL dox. Subsequently, 50% of the medium was replaced with fresh neuronal medium (lacking dox and Ara-C) once every other day until the neurons were fixed or harvested at day 21.

#### ii) shRNA treated neurons (Fig. 1H,I)

On day −1 NPCs were dissociated with Accutase Cell Detachment Solution (Innovative Cell Technologies, # AT-104) for 5min at 37°C, counted and seeded at a density of at 5⨯10^5^ cells/well on Matrigel coated 24-well plates in hNPC media (DMEM/F12 (Life Technologies #10565), 1x N-2 (Life Technologies #17502-048), 1x B-27-RA (Life Technologies #12587-010), 20 ng/mL FGF2 (Life Technologies)) on Matrigel (Corning, #354230). On day 0, cells were transduced with rtTA (Addgene 20342) and *NGN2* (Addgene 99378) lentiviruses as well as desired shRNA viruses in NPC media containing 10 μM Thiazovivin (Millipore, #S1459) and spinfected (centrifuged for 1 hour at 1000g). On day 1, media was replaced and dox was added with 1ug/mL working concentration. On day 2, transduced hNPCs were treated with corresponding antibiotics to the lentiviruses (1 μg/mL puromycin for *shRNA*, 1 mg/mL G-418 for *NGN2*-Neo). On day 4, medium was switched to Brainphys neuron medium (Brainphys (STEMCELL, # 05790), 1% N-2, 2% B-27-RA, 1 μg/mL Natural Mouse Laminin (Thermofisher, # 23017015), 10 ng/mL BDNF (R&D, #248), 10 ng/mL GDNF (R&D, #212), 500 μg/mL Dibutyryl cyclic-AMP (Sigma, #D0627), 200 nM L-ascorbic acid (Sigma, # A4403) and 1 μg/mL dox. Medium was replaced every second day until SARS-CoV-2 infection on day 7.

#### iii) rs4702 neurons (Fig. 5)

On day 0, hiPSCs were transduced with rtTA (Addgene 20342) and NGN2 (Addgene 99378) lentiviruses. hiPSCs were infected in a conical tube in low volume StemFlex media and high virus concentration and subsequently seeded onto 6-wells at 1⨯10^5^ cells/well and spinfected. On day 1, Medium was switched to non-viral neurobasal neuron medium (Neurobasal (Thermofisher Scientific, #21103049), 1x N-2 (Life Technologies #17502-048), 1x B-27-RA (Life Technologies #12587-010), 1 μg/mL Natural Mouse Laminin (Life Technologies), 20 ng/mL BDNF (Peprotech #450-02), 20 ng/mL GDNF (Peprotech #450-10), 500 μg/mL Dibutyryl cyclic-AMP (Sigma #D0627), 200 nM L-ascorbic acid (Sigma #A0278)) and Doxycycline (dox) was added to each well at a final concentration of 1 μg/mL. On day 2, transduced hiPSCs were treated with corresponding antibiotics to the lentiviruses (1 mg/mL G-418). On day 4, medium was replaced including 1 μg/mL dox and 4 μM cytosine arabinoside (Ara-C) to reduce the proliferation of non-neuronal cells along with antibiotic withdrawal. On day 6 medium was replaced (including 1 μg/mL dox). At day 7, neurons were infected.

### Differentiation of lung alveolospheres (Jacob et al., 2019)

Alveolospheres, composed of distal lung epithelial cells, were generated from hiPSCs cultured in mTeSR Plus medium (STEMCELLTechnologies, #05825) through a sequential directed differentiation protocol (Jacob et al., 2019). Developmental lung epithelial lineage differentiation was initiated by dissociating hiPSCs with Accutase Cell Detachment Solution (Innovative Cell Technologies, # AT-104) and seeding 2⨯10^6^ cells onto one well of a matrigel-coated 6-well plate in mTeSR Plus medium with 10 μM THX. For the next three days, cells were patterned into definitive endoderm using the StemDiff Definitive Endoderm kit (STEMCELL Technologies, #05110). On day 3, cells were dissociated with Gentle Cell Dissociation Reagent (STEMCELL Technologies 07174 and passaged 1:4 (C1) or 1:6 (C2) and seeded in DS/SB Media (cSFDM Base (Jacob et al., 2019), 10 μM SB43152 (Tocris 1614), 2 μM Dorsomorphin (Stemgent 04-0024)) plus 10 μM THX. Followed by patterning into anterior foregut endoderm (until day 6) using inhibition of BMP/TGFB signaling. Specification of lung lineage was achieved by using three factors (CHIR99021, BMP4, and retinoic acid) to produce primordial lung progenitors which were enriched by sorting using the cell-surface markers CD47hi/CD26lo (Jacob et al., 2019; McCauley et al., 2018) and then plated in Matrigel for 3D culture on day 15. These progenitors were then differentiated into distal lung epithelium using CHIR99021, keratinocyte growth factor, dexamethasone, cyclic AMP, and 3-isobutyl-1-methyxanthine). After day 25, monolayered epithelial spheres (“alveolospheres”) emerged, and were infected and harvested as specified, typically ∼day 30.

For shRNA experiments, lung progenitors were collected, dissociated and seeded as single cells in the presence of ROCK inhibitor THX at day 14. The following day, these cells were infected with lentiviral shRNAs against SARS-CoV-2 host genes, followed by 2 days of puromycin selection. One day after withdrawal of antibiotic selection for dissociated alveolar cells (or media change for control 3D alveolospheres), cells were infected with SARS-CoV-2 (MOI 0.1-0.5) on day 33 and harvested 24 hours later.

### Differentiation of intestinal organoids

(Koike et al., 2019; Zhang et al., 2018) Human posterior gut endoderm can be generated from hiPSCs through a stepwise differentiation protocol adapted from (Koike et al., 2019; Zhang et al., 2018). The first step is patterning to definitive endoderm by initial treatment with activin A and Wnt3a (2 days), followed by BMP4, FGF2, VEGF, and activin A treatment (through day 6). Definitive endoderm is re-plated and matured with FGF2, A83-01 and CHIR99021 into CDX2+ posterior gut endoderm progenitor cells. Gut progenitors can be expanded (FGF2, VEGF, EGF, A83-01 and CHIR99021) and differentiated into gut organoids, simply by treating with CHIR 99021 and FGF4 for 3 days until dissociated intestinal cell sheets started budding off and formed floating hindgut spheroids. After 7 additional days embedded in Matrigel, spheroids differentiate and protrude into the lumen. Hindgut organoids comprise a polarized epithelium that includes absorptive enterocytes (VILLIN+) and hindgut epithelial cells (E-CAD+). On day 20, intestinal organoids were collected and dissociated and seeded as single cells. The next day, cells were infected with shRNA viruses against SARS-CoV-2 receptor targets, followed by 2 days of puromycin selection. One day after selection withdrawal for dissociated intestinal cells and regular feeding for 3D intestinal organoids, cells were infected with SARS-CoV-2 (MOI 0.05-1) on day 25 or day 50 and harvested 24 or 48 hours later.

### Lentivirus generation

Third-generation VSV.G pseudotyped HIV-1 lentiviruses were produced by polyethylenimine (PEI, Polysciences #23966-2)-transfection of HEK293T cells and packaged with VSVG-coats using established methods. Used shRNA against *ACE2* (SHCLNG-NM_021804), *CD147* (SHCLNG-NM_001728), *TMPRSS2* (SHCLNG-NM_005656) were obtained from Sigma.

### SARS-CoV-2 virus propagation and infections

SARS-related coronavirus 2 (SARS-CoV-2), isolate USA-WA1/2020 (NR-52281) was deposited by the Center for Disease Control and Prevention and obtained through BEI Resources, NIAID, NIH. SARS-CoV-2 was propagated in Vero E6 cells in DMEM supplemented with 2% FBS, 4.5 g/L D-glucose, 4 mM L-glutamine, 10 mM Non-Essential Amino Acids, 1 mM Sodium Pyruvate and 10 mM HEPES. Virus stock was filtered by centrifugation using Amicon Ultra-15 Centrifugal filter unit (Sigma, Cat # UFC910096) and resuspended in viral propagation media. All infections were performed with either passage 3 or 4 SARS-CoV-2. Infectious titers of SARS-CoV-2 were determined by plaque assay in Vero E6 cells in Minimum Essential Media supplemented with 4 mM Lglutamine, 0.2% BSA, 10 mM HEPES and 0.12% NaHCO3 and 0.7% Oxoid agar (Cat #OXLP0028B). All SARS-CoV-2 infections were performed in the CDC/USDA-approved BSL-3 facility of the Global Health and Emerging Pathogens Institute at the Icahn School of Medicine at Mount Sinai in accordance with institutional biosafety requirements.

### RNA-seq of SARS-CoV-2 infected hamster brains

3-5-week-old male Golden Syrian hamsters (*Mesocricetus auratus*) were obtained from Jackson Laboratories. Hamsters were acclimated to the CDC/USDA-approved BSL-3 facility of the Global Health and Emerging Pathogens Institute at the Icahn School of Medicine at Mount Sinai for 2-4 days. Before intranasal infection, hamsters were anesthetized by intraperitoneal injection with 200 μl of a ketamine HCl/xyalzine (4:1) solution. Inoculum at the referenced doses (high and low, 100pfu and 10,000 pfu, respectively) were resuspended in PBS to a total volume of 100 μL. All hamsters analyzed by RNA-seq were perfused with PBS before brains were harvested. The brain was cut in half longitudinally after removing hindbrain and half was used for RNA extraction and analysis. Brain tissues were homogenized for 40 seconds at 6.00 m/s for 2 cycles (MP Biomedicals, Cat#: SKU 116005500) in Lysing Matrix A homogenization tubes (MP Biomedicals, Cat#6910-100). All tissues for RNA-seq analysis were homogenized directly in TRIzol (Invitrogen, Cat#15596026). RNA was isolated by phenol/chloroform extraction according to manufacturer’s instructions.

RNA-seq libraries were prepared using the TruSeq Stranded mRNA Library Prep kit (Illumina) according to the manufacturer’s instructions. cDNA libraries were sequenced and single-end sequencing reads were aligned using an Illumina NextSeq 500 platform. Reads were aligned to *Mesocricetus auratus* Ensembl MesAur1.0 using STAR aligner, and mappid using CustomDESeq2.

RNA sequencing analysis was performed in R v.3.6.0. Differential expression analysis was performed using edgeR (v3.26.8) (Robinson et al., 2010), limma (v3.40.6) (Ritchie et al., 2015) and Glimma (v1.12.0) (Su et al., 2017) packages. Raw read counts were transformed into log2 CPM using the edgeR package. The limma voom function was used to compute the weights for heteroscedasticity adjustment by estimating the mean variance trend for log2 counts. Linear models were fitted to the expression values of each gene using the lmFit function. Empirical Bayesian moderation was applied using the eBayes function to obtain more precise estimates of gene-wise variability. *P* values were adjusted for multiple hypotheses testing using FDR estimation; DEGs were determined as those with an estimated FDR ≤ 5%. Enrichment analysis was performed using WebGestaltR (0.4.4) using custom neural gene sets of interest (Schrode et al., 2019) and interferon gene sets (Blanco-Melo et al., 2020). Genes of interest were ranked by −log10(*P*) and enrichment was performed against a background of all expressed genes.

### Molecular and biochemical analysis

#### i. Real time-quantitative PCR

Cell were harvested with TRIzol and total RNA extraction was carried out following the manufacturer’s instructions. Quantitative transcript analysis was performed using a QuantStudio 7 Flex Real-Time PCR System with the Power SYBR Green RNA-to-Ct Real-Time qPCR Kit (all ThermoFisher). Total RNA template (25 ng per reaction) was added to the PCR mix, including primers. qPCR conditions were as follows; 48°C for 15 min, 95°C for 10 min followed by 45 cycles (95°C for 15 s, 60°C for 60 s). All qPCR data is collected from at least 3 independent biological replicates of one experiment. If not otherwise stated, obtained Ct values were normalized against 18S one super-control (C1 alveolosphere MOI 0.1) to ensure comparability across plates, experiments and cell types (except for shRNA experiments, which were normalized to either infected or uninfected scrambled control). Data analyses were performed using GraphPad PRISM 8 software.

Primers were used as follows

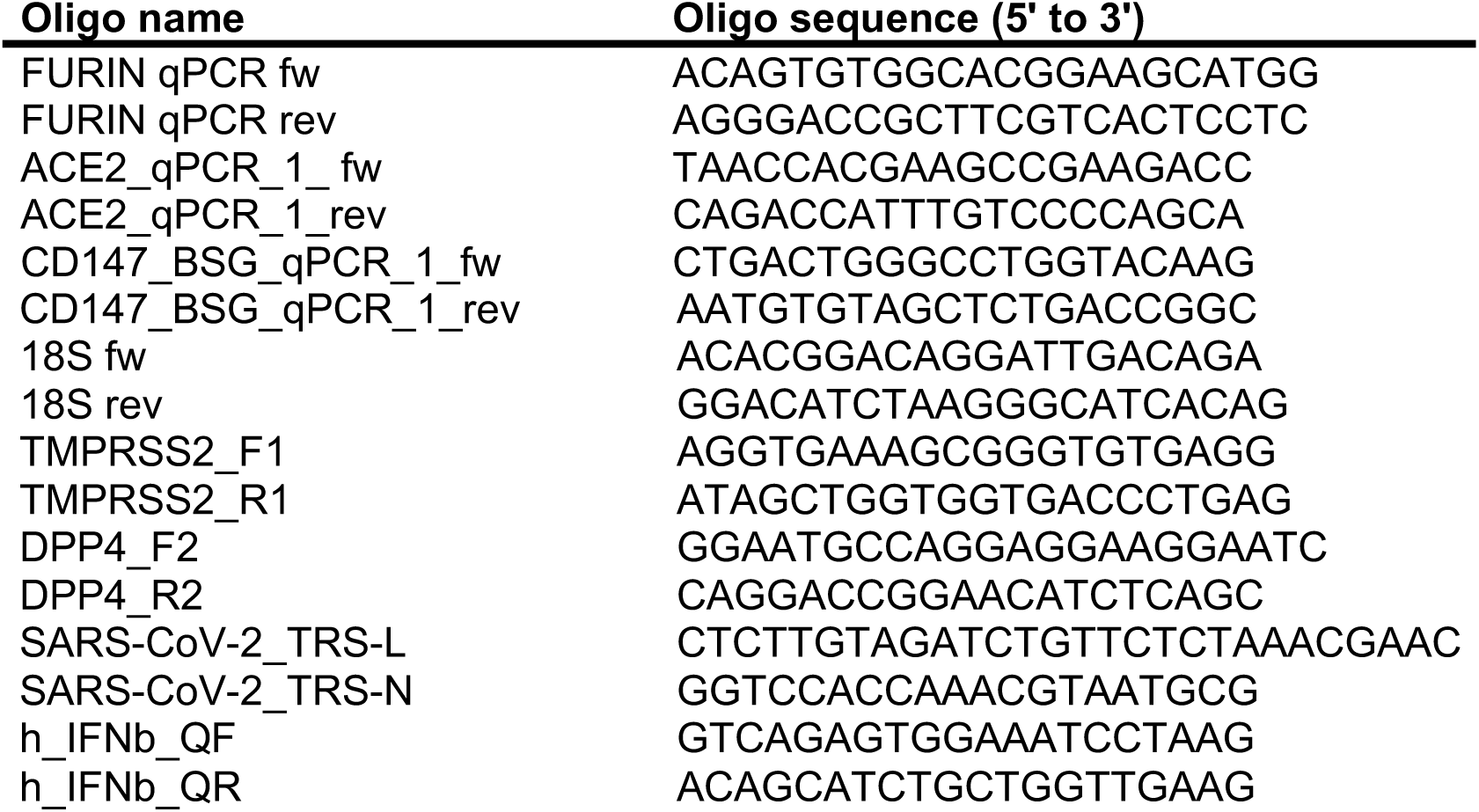

#### ii. Fluorescent in situ hybridization (FISH)

RNA FISH was performed on post-mortem brain (74-year-old male donor, post-mortem interval 87 hours) using the RNAscope Multiplex Fluorescent v2 protocol (Advanced Cell Diagnostics) after fixation with 4% formaldehyde for 6 hours at 4°C, followed by incubation in 30% sucrose overnight. Immediately after the *in situ* labeling was completed, endogenous background was reduced using TrueBlack Lipofuscin Autofluorescence Quencher (Biotium). Sections were Mounted with Vectashield antifade medium and images were acquired on a Zeiss LSM780 inverted confocal microscope.

#### iii. Immunostaining and microscopy

Cells were washed with ice-cold PBS and fixed with 4% PFA solution at pH 7.4 overnight at 4**°**C. Then, fixative solution was replaced with PBS (Ca/Mg++). Alveolar and intestinal cells were permeabilized and blocked with 0.1% Triton-X in PBS (Ca/Mg++) + 2% donkey serum for 1 hour at room temperature. Neurons were permeabilized and blocked with 0.05% Tween in PBS (Ca/Mg++) + 2% donkey serum for 1 hour at room temperature. The blocking solution was aspirated and replaced with the same solution with primary antibodies. Mouse monoclonal anti-SARS-CoV-2 Nucleocapsid [1C7C7] protein (1:1000, a kind gift by Dr. T. Moran, Center for Therapeutic Antibody Discovery at the Icahn School of Medicine at Mount Sinai) and rabbit anti-ß-III-tubulin (Convence, PRB-435P, 1:2000) or goat anti-EpCAM, (R&D Biosystems #AF960, 1:500) overnight at 4°C. Cells were washed 3x for 5 min with PBS (Ca/Mg++) and subsequently incubated with secondary antibodies, prepared in blocking solution, for 1 hour at room temperature, followed by 300 nM DAPI staining for 3-4 min and washed 2x for 5 min with PBS (Ca/Mg++).

Organoids were fixed with 4% PFA and incubated in 20% sucrose overnight, then frozen in OCT. Frozen organoids were cryosectioned to 10um sections and mounted on VWR Superfrost Plus slides. Slides were washed and permeabilized with 0.01% Triton-X PBS and incubated with blocking solution (2% donkey serum in PBS) for 1 hour. Then slides were incubated with primary antibody (EpCAM, 1:50, R&D Biosystems #AF960; Villin, Abcam, ab130751, COV2-NP 1:100, ACE2 1:100) overnight. Slides were then incubated with corresponding anti-mouse, anti-rabbit, and anti-goat secondary antibodies (1:200, 2 hrs) and DAPI (1:2000, 10 mins) and mounted with ImmuMount.

Cells and organoids were imaged with a Zeiss LSM 780, Nikon C2 confocal microscope or ThermoFisher HCS CX7 microscope. The images were quantified using the CellProfiler software. Data points represent at least 100 cells or one 96-well each.

#### iv. Immunoblot

For the Western Blot Analysis of COVID-19 receptors in human brain, 100 mg (1-volume) of brain tissue was Dounce homogenized in 1 mL (10 volumes) of ice-cold protein extraction buffer (50 mM-Tris-HCl, pH 8; 20 mM-NaCl; 2 mM-MgCl2; 4 M-Urea, 0.35% Triton-X100; 0.35% Sodium deoxycholate; 0.05% SDS; supplemented with protease and phosphatase inhibitors). Benzonase was added to eliminate viscosity of the sample for 10 min on ice, then sonicated and centrifuged at 20,817 x g for 15 minutes at 4°C. Clarified total lysates were quantified using a BCA Protein Assay kit (Pierce). Western blot analysis was performed using 100 μg of total protein using anti-ACE2 (Abcam/ab239924/EPR4435(2)), anti-CD147 (Abcam/ab666/MEM-M6/1), anti-FURIN (Abcam/ab183495/EPR14674) or anti-β-ACTIN (Cell Signaling/#4970S/ 13E5) primary antibodies (1:5000, 1-hr) followed by autoradiographic detection using corresponding anti-mouse (Cell Signaling/#7076S) or anti-rabbit (Cell Signaling/#7074S) secondary antibodies conjugated with HRP (1:5000, 1-hr). For cell cultures, weight of the harvested cell pellet was used to prepare the extracts and followed the rest of the protocol as described above.

#### V. Data analysis

Data from all phenotypic assays above were first organized in a Microsoft Excel spreadsheet and analyzed using GraphPad PRISM 8 software or R. For qPCR data analysis, values are expressed as mean ± SEM. Statistical significance was tested using either one-sided student T-test or two-way ANOVA with Tukey’s post-hoc test for comparison of all sample means.

## STATEMENT OF ETHICS

Ethical approval was not required because the hiPSC lines, lacking association with any identifying information and widely accessible from a public repository, are thus not considered to be human subjects research.

## AUTHOR CONTRIBUTIONS

KJB, SA and BT conceptualized this collaborative approach. KJB, SA, BT, KD, DAH. contributed to experimental design and wrote the manuscript. KD coordinated all hiPSC-based studies and led the neuron differentiation, with BK leading lung differentiation, CS leading intestinal differentiations. DK provided critical expertise in lung organoid differentiation and KI and TT provided not just expertise in intestinal organoids, but also frozen differentiated intestinal organoids. DAH coordinated all SARS-CoV-2 infections, assisted by RM and KD. qPCR experiments were supported by CO, MI, BK and MFG. Post-mortem brain studies by BJ, CP, JC, and SA. High-content imaging and viral production was carried out by KD and assisted by PJMD. Additional neuronal inductions and organoid sectioning and staining supported by AM and MBF. Bioinformatics of hiPSC neurons by SP and hamster brains by CS.

## DATA AVAILABILITY

All donor-derived and CRISPR-edited hiPSCs are available at the NIMH repository at RUCDR.

---

**Supplemental Figure 1 (related to Figure 1).**
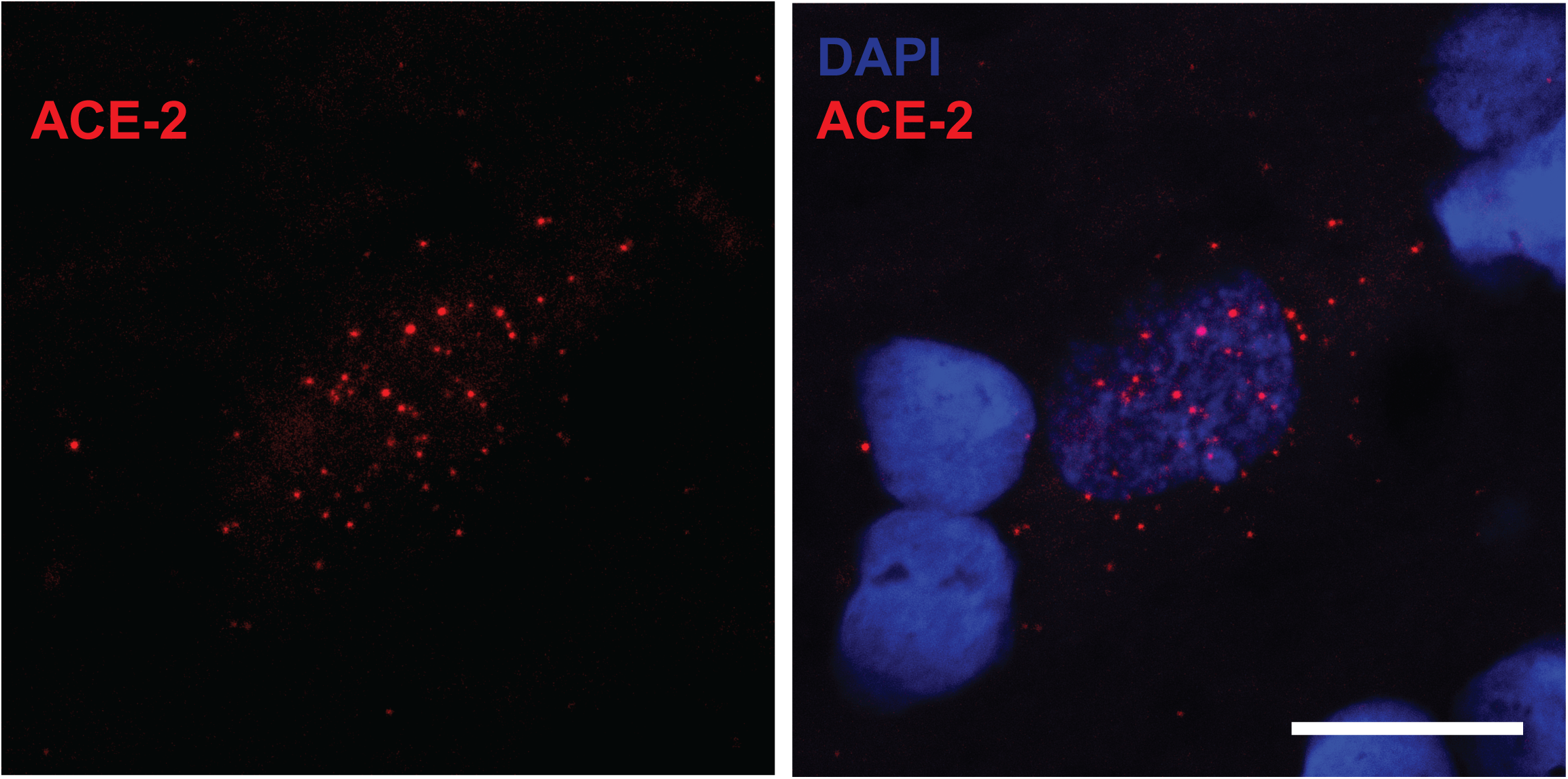
*ACE2* in situ hybridization RNA staining over post-mortem human ventral midbrain cell with neuronal nucleus-like morphology in DAPI stain. Scale bar: 20μm.

**Supplemental Table 1.** Excess of nominally significant GWAS variants within the SARS-CoV-2 host gene cis-regions when comparing individuals with COVID-19 to the general population.

| Gene | Study name | Phenotype | Case number | Control number | N SNPs in cis-region (+/- 1e06) | N nominal (p<0.05) | % reaching nominal significance | binomial p value |
| --- | --- | --- | --- | --- | --- | --- | --- | --- |
| *ACE2* | COVID19_HGI_ANA_B2 | hospitalized covid vs. population | 3199 | 897488 | 4839 | 480 | 9.919405 | 3.28E-44 |
| *ACE2* | COVID19_HGI_ANA_C2 | covid vs. population | 6696 | 1073072 | 5281 | 497 | 9.411096 | 1.02E-39 |
| *CD147* | COVID19_HGI_ANA_D1 | predicted covid from self-reported symptoms vs. predicted or self-reported non-covid | 1865 | 29174 | 6456 | 584 | 9.045849 | 2.54E-41 |
| *FURIN* | COVID19_HGI_ANA_B2 | hospitalized covid vs. population | 3199 | 897488 | 11556 | 869 | 7.519903 | 3.57E-31 |
| *TMPRSS2* | COVID19_HGI_ANA_B2 | hospitalized covid vs. population | 3199 | 897488 | 14047 | 919 | 6.542322 | 1.00E-15 |
| *TMPRSS2* | COVID19_HGI_ANA_C1 | covid vs. lab/self-reported negative | 3523 | 36634 | 20141 | 1104 | 5.481356 | 0.00201 |
| *FURIN* | COVID19_HGI_ANA_C1 | covid vs. lab/self-reported negative | 3523 | 36634 | 17346 | 921 | 5.309581 | 6.00E-02 |
| *TMPRSS2* | COVID19_HGI_ANA_C2 | covid vs. population | 6696 | 1073072 |  | 21814 | 4.776749 |  |
| *TMPRSS2* | COVID19_HGI_ANA_D1 | predicted covid from self-reported symptoms vs. predicted or self-reported non-covid | 1865 | 29174 |  | 12030 | 4.763092 |  |
| *CD147* | COVID19_HGI_ANA_C2 | covid vs. population | 6696 | 1073072 |  | 14419 | 4.722935 |  |
| *CD147* | COVID19_HGI_ANA_B2 | hospitalized covid vs. population | 3199 | 897488 |  | 9554 | 4.406531 |  |
| *CD147* | COVID19_HGI_ANA_B2 | hospitalized covid vs. population | 3199 | 897488 |  | 9554 | 4.406531 |  |
| *CD147* | COVID19_HGI_ANA_C1 | covid vs. lab/self-reported negative | 3523 | 36634 |  | 13358 | 3.960174 |  |
| *FURIN* | COVID19_HGI_ANA_C2 | covid vs. population | 6696 | 1073072 |  | 19136 | 3.88796 |  |
| *ACE2* | COVID19_HGI_ANA_D1 | predicted covid from self-reported symptoms vs. predicted or self-reported non-covid | 1865 | 29174 |  | 532 | 3.383459 |  |
| *ACE2* | COVID19_HGI_ANA_C1 | covid vs. lab/self-reported negative | 3523 | 36634 |  | 4098 | 3.29429 |  |
| *FURIN* | COVID19_HGI_ANA_D1 | predicted covid from self-reported symptoms vs. predicted or self-reported non-covid | 1865 | 29174 |  | 9432 | 3.042833 |  |

## REFERENCES

Abo, K.M., Ma, L., Matte, T., Huang, J., Alysandratos, K.D., Werder, R.B., Mithal, A., Beermann, M.L., Lindstrom-Vautrin, J., Mostoslavsky, G., et al. (2020). Human iPSC-derived alveolar and airway epithelial cells can be cultured at air-liquid interface and express SARS-CoV-2 host factors. bioRxiv.

Barretto, N., Zhang, H., Powell, S.K., Fernando, M.B., Zhang, S., Flaherty, E.K., Ho, S.M., Slesinger, P.A., Duan, J., and Brennand, K.J. (2020). ASCL1- and DLX2-induced GABAergic neurons from hiPSC-derived NPCs. J Neurosci Methods 334, 108548.

Belouzard, S., Chu, V.C., and Whittaker, G.R. (2009). Activation of the SARS coronavirus spike protein via sequential proteolytic cleavage at two distinct sites. Proc Natl Acad Sci U S A 106, 5871–5876.

Blanco-Melo, D., Nilsson-Payant, B.E., Liu, W.C., Uhl, S., Hoagland, D., Moller, R., Jordan, T.X., Oishi, K., Panis, M., Sachs, D., et al. (2020). Imbalanced Host Response to SARS-CoV-2 Drives Development of COVID-19. Cell 181, 1036–1045 e1039.

Chen, X., Laurent, S., Onur, O.A., Kleineberg, N.N., Fink, G.R., Schweitzer, F., and Warnke, C. (2020). A systematic review of neurological symptoms and complications of COVID-19. J Neurol.

Chu, C.M., Poon, L.L., Cheng, V.C., Chan, K.S., Hung, I.F., Wong, M.M., Chan, K.H., Leung, W.S., Tang, B.S., Chan, V.L., et al. (2004). Initial viral load and the outcomes of SARS. CMAJ 171, 1349–1352.

Ciancanelli, M.J., Huang, S.X., Luthra, P., Garner, H., Itan, Y., Volpi, S., Lafaille, F.G., Trouillet, C., Schmolke, M., Albrecht, R.A., et al. (2015). Infectious disease. Life-threatening influenza and impaired interferon amplification in human IRF7 deficiency. Science 348, 448–453.

Coutard, B., Valle, C., de Lamballerie, X., Canard, B., Seidah, N.G., and Decroly, E. (2020). The spike glycoprotein of the new coronavirus 2019-nCoV contains a furin-like cleavage site absent in CoV of the same clade. Antiviral Res 176, 104742.

Cuomo, A.S.E., Seaton, D.D., McCarthy, D.J., Martinez, I., Bonder, M.J., Garcia-Bernardo, J., Amatya, S., Madrigal, P., Isaacson, A., Buettner, F., et al. (2020). Single-cell RNA-sequencing of differentiating iPS cells reveals dynamic genetic effects on gene expression. Nat Commun 11, 810.

Das, R., and Ghate, S.D. (2020). Investigating the likely association between genetic ancestry and COVID-19 manifestation. medRxiv, 2020.2004.2005.20054627.

Desforges, M., Le Coupanec, A., Brison, E., Meessen-Pinard, M., and Talbot, P.J. (2014). Neuroinvasive and neurotropic human respiratory coronaviruses: potential neurovirulent agents in humans. Adv Exp Med Biol 807, 75–96.

Dixit, A., Parnas, O., Li, B., Chen, J., Fulco, C.P., Jerby-Arnon, L., Marjanovic, N.D., Dionne, D., Burks, T., Raychowdhury, R., et al. (2016). Perturb-Seq: Dissecting Molecular Circuits with Scalable Single-Cell RNA Profiling of Pooled Genetic Screens. Cell 167, 1853–1866 e1817.

Dutta, M., Dutta, P., Medhi, S., Borkakoty, B., and Biswas, D. (2018). Polymorphism of HLA class I and class II alleles in influenza A(H1N1)pdm09 virus infected population of Assam, Northeast India. J Med Virol 90, 854–860.

Ellinghaus, D., Degenhardt, F., Bujanda, L., Buti, M., Albillos, A., Invernizzi, P., Fernandez, J., Prati, D., Baselli, G., Asselta, R., et al. (2020). The ABO blood group locus and a chromosome 3 gene cluster associate with SARS-CoV-2 respiratory failure in an Italian-Spanish genome-wide association analysis. medRxiv, 2020.2005.2031.20114991.

Everitt, A.R., Clare, S., Pertel, T., John, S.P., Wash, R.S., Smith, S.E., Chin, C.R., Feeley, E.M., Sims, J.S., Adams, D.J., et al. (2012). IFITM3 restricts the morbidity and mortality associated with influenza. Nature 484, 519–523.

Forrest, M.P., Zhang, H., Moy, W., McGowan, H., Leites, C., Dionisio, L.E., Xu, Z., Shi, J., Sanders, A.R., Greenleaf, W.J., et al. (2017). Open Chromatin Profiling in hiPSC-Derived Neurons Prioritizes Functional Noncoding Psychiatric Risk Variants and Highlights Neurodevelopmental Loci. Cell stem cell.

Franzen, O., Gan, L.M., and Bjorkegren, J.L.M. (2019). PanglaoDB: a web server for exploration of mouse and human single-cell RNA sequencing data. Database (Oxford) 2019.

Fromer, M., Roussos, P., Sieberts, S.K., Johnson, J.S., Kavanagh, D.H., Perumal, T.M., Ruderfer, D.M., Oh, E.C., Topol, A., Shah, H.R., et al. (2016). Gene expression elucidates functional impact of polygenic risk for schizophrenia. Nat Neurosci 19, 1442–1453.

Gasperini, M., Hill, A.J., McFaline-Figueroa, J.L., Martin, B., Kim, S., Zhang, M.D., Jackson, D., Leith, A., Schreiber, J., Noble, W.S., et al. (2019). A Genome-wide Framework for Mapping Gene Regulation via Cellular Genetic Screens. Cell 176, 377–390 e319.

Gordon, D.E., Jang, G.M., Bouhaddou, M., Xu, J., Obernier, K., White, K.M., O’Meara, M.J., Rezelj, V.V., Guo, J.Z., Swaney, D.L., et al. (2020). A SARS-CoV-2 protein interaction map reveals targets for drug repurposing. Nature.

GTEx Consortium, Battle, A., Brown, C.D., Engelhardt, B.E., and Montgomery, S.B. (2017). Genetic effects on gene expression across human tissues. Nature 550, 204–213.

Guan, W.J., Ni, Z.Y., Hu, Y., Liang, W.H., Ou, C.Q., He, J.X., Liu, L., Shan, H., Lei, C.L., Hui, D.S.C., et al. (2020). Clinical Characteristics of Coronavirus Disease 2019 in China. N Engl J Med.

Han, Y., Yang, L., Duan, X., Duan, F., Nilsson-Payant, B.E., Yaron, T.M., Wang, P., Tang, X., Zhang, T., Zhao, Z., et al. (2020). Identification of Candidate COVID-19 Therapeutics using hPSC-derived Lung Organoids. bioRxiv.

Hellberg, A., Poole, J., and Olsson, M.L. (2002). Molecular basis of the globoside-deficient P(k) blood group phenotype. Identification of four inactivating mutations in the UDP-N-acetylgalactosamine: globotriaosylceramide 3-beta-N-acetylgalactosaminyltransferase gene. J Biol Chem 277, 29455–29459.

Helms, J., Kremer, S., Merdji, H., Clere-Jehl, R., Schenck, M., Kummerlen, C., Collange, O., Boulay, C., Fafi-Kremer, S., Ohana, M., et al. (2020). Neurologic Features in Severe SARS-CoV-2 Infection. N Engl J Med 382, 2268–2270.

Ho, S.M., Hartley, B.J., Tcw, J., Beaumont, M., Stafford, K., Slesinger, P.A., and Brennand, K.J. (2016). Rapid Ngn2-induction of excitatory neurons from hiPSC-derived neural progenitor cells. Methods 101, 113–124.

Hoffman, G.E., Hartley, B.J., Flaherty, E., Ladran, I., Gochman, P., Ruderfer, D.M., Stahl, E.A., Rapoport, J., Sklar, P., and Brennand, K.J. (2017). Transcriptional signatures of schizophrenia in hiPSC-derived NPCs and neurons are concordant with post-mortem adult brains. Nat Commun 8, 2225.

Hoffmann, M., Kleine-Weber, H., Schroeder, S., Kruger, N., Herrler, T., Erichsen, S., Schiergens, T.S., Herrler, G., Wu, N.H., Nitsche, A., et al. (2020). SARS-CoV-2 Cell Entry Depends on ACE2 and TMPRSS2 and Is Blocked by a Clinically Proven Protease Inhibitor. Cell.

Hou, Y., Liang, W., Zhang, J., Li, Q., Ou, H., Wang, Z., Li, S., Huang, X., and Zhao, C. (2018). Schizophrenia-associated rs4702 G allele-specific downregulation of FURIN expression by miR-338-3p reduces BDNF production. Schizophr Res 199, 176–180.

Huang, J., Hume, A.J., Abo, K.M., Werder, R.B., Villacorta-Martin, C., Alysandratos, K.D., Beermann, M.L., Simone-Roach, C., Olejnik, J., Suder, E.L., et al. (2020). SARS-CoV-2 Infection of Pluripotent Stem Cell-derived Human Lung Alveolar Type 2 Cells Elicits a Rapid Epithelial-Intrinsic Inflammatory Response. bioRxiv.

Imai, M., Iwatsuki-Horimoto, K., Hatta, M., Loeber, S., Halfmann, P.J., Nakajima, N., Watanabe, T., Ujie, M., Takahashi, K., Ito, M., et al. (2020). Syrian hamsters as a small animal model for SARS-CoV-2 infection and countermeasure development. Proc Natl Acad Sci U S A 117, 16587–16595.

Initiative, C.-H.G. (2020). The COVID-19 Host Genetics Initiative, a global initiative to elucidate the role of host genetic factors in susceptibility and severity of the SARS-CoV-2 virus pandemic. Eur J Hum Genet 28, 715–718.

Jacob, A., Vedaie, M., Roberts, D.A., Thomas, D.C., Villacorta-Martin, C., Alysandratos, K.D., Hawkins, F., and Kotton, D.N. (2019). Derivation of self-renewing lung alveolar epithelial type II cells from human pluripotent stem cells. Nat Protoc 14, 3303–3332.

Jacob, F., Pather, S.R., Huang, W.-K., Wong, S.Z.H., Zhou, H., Zhang, F., Cubitt, B., Chen, C.Z., Xu, M., Pradhan, M., et al. (2020). Human Pluripotent Stem Cell-Derived Neural Cells and Brain Organoids Reveal SARS-CoV-2 Neurotropism. bioRxiv, 2020.2007.2028.225151.

Jerber, J., Seaton, D., Cuomo, A., Kumasaka, N., Haldane, J., Steer, J., Patel, M., Pearce, D., Andersson, M., Bonder, M., et al. (2020). Population-scale single-cell RNA-seq profiling across dopaminergic neuron differentiation. bioRxiv, 2020.2005.2021.103820.

Kachuri, L., Francis, S.S., Morrison, M., Wendt, G., Bossé, Y., Cavazos, T.B., Rashkin, S.R., Ziv, E., and Witte, J.S. (2020). The landscape of host genetic factors involved in immune response to common viral infections. medRxiv, 2020.2005.2001.20088054.

Karczewski, K.J., Francioli, L.C., Tiao, G., Cummings, B.B., Alföldi, J., Wang, Q., Collins, R.L., Laricchia, K.M., Ganna, A., Birnbaum, D.P., et al. (2020). The mutational constraint spectrum quantified from variation in 141,456 humans. bioRxiv Nature (in press), 531210.

Kenney, A.D., Dowdle, J.A., Bozzacco, L., McMichael, T.M., St Gelais, C., Panfil, A.R., Sun, Y., Schlesinger, L.S., Anderson, M.Z., Green, P.L., et al. (2017). Human Genetic Determinants of Viral Diseases. Annu Rev Genet 51, 241–263.

Koike, H., Iwasawa, K., Ouchi, R., Maezawa, M., Giesbrecht, K., Saiki, N., Ferguson, A., Kimura, M., Thompson, W.L., Wells, J.M., et al. (2019). Modelling human hepato-biliary-pancreatic organogenesis from the foregut-midgut boundary. Nature 574, 112–116.

Konno, Y., Kimura, I., Uriu, K., Fukushi, M., Irie, T., Koyanagi, Y., Sauter, D., Gifford, R.J., Consortium, U.-C., Nakagawa, S., et al. (2020). SARS-CoV-2 ORF3b Is a Potent Interferon Antagonist Whose Activity Is Increased by a Naturally Occurring Elongation Variant. Cell reports, 108185.

Lamers, M.M., Beumer, J., van der Vaart, J., Knoops, K., Puschhof, J., Breugem, T.I., Ravelli, R.B.G., Paul van Schayck, J., Mykytyn, A.Z., Duimel, H.Q., et al. (2020). SARS-CoV-2 productively infects human gut enterocytes. Science 369, 50–54.

Lei, X., Dong, X., Ma, R., Wang, W., Xiao, X., Tian, Z., Wang, C., Wang, Y., Li, L., Ren, L., et al. (2020). Activation and evasion of type I interferon responses by SARS-CoV-2. Nat Commun 11, 3810.

Li, Y.C., Bai, W.Z., and Hashikawa, T. (2020). The neuroinvasive potential of SARS-CoV2 may play a role in the respiratory failure of COVID-19 patients. J Med Virol.

Lindesmith, L., Moe, C., Marionneau, S., Ruvoen, N., Jiang, X., Lindblad, L., Stewart, P., LePendu, J., and Baric, R. (2003). Human susceptibility and resistance to Norwalk virus infection. Nat Med 9, 548–553.

Ludvigsson, J.F. (2020). Systematic review of COVID-19 in children shows milder cases and a better prognosis than adults. Acta Paediatr 109, 1088–1095.

Mao, L., Wang, M., Chen, S., He, Q., Chang, J., Hong, C., Zhou, Y., Wang, D., Li, Y., Jin, H., et al. (2020). Neurological Manifestations of Hospitalized Patients with COVID-19 in Wuhan, China: a retrospective case series study. medRxiv, 2020.2002.2022.20026500.

McCauley, K.B., Hawkins, F., and Kotton, D.N. (2018). Derivation of Epithelial-Only Airway Organoids from Human Pluripotent Stem Cells. Curr Protoc Stem Cell Biol 45, e51.

McLoughlin, B.C., Miles, A., Webb, T.E., Knopp, P., Eyres, C., Fabbri, A., Humphries, F., and Davis, D. (2020). Functional and cognitive outcomes after COVID-19 delirium. Eur Geriatr Med.

Mimitou, E.P., Cheng, A., Montalbano, A., Hao, S., Stoeckius, M., Legut, M., Roush, T., Herrera, A., Papalexi, E., Ouyang, Z., et al. (2019). Multiplexed detection of proteins, transcriptomes, clonotypes and CRISPR perturbations in single cells. Nat Methods 16, 409–412.

Mitchell, J.M., Nemesh, J., Ghosh, S., Handsaker, R.E., Mello, C., Meyer, D., Raghunathan, K., de Rivera, H., Tegtmeyer, M., Hawes, D., et al. (2020). Mapping genetic effects on cellular phenotypes with “cell villages”. bioRxiv, 2020.2006.2029.174383.

Neavin, D., Nguyen, Q., Daniszewski, M.S., Liang, H.H., Chiu, H.S., Senabouth, A., Lukowski, S.W., Crombie, D.E., Lidgerwood, G.E., Hernández, D., et al. (2020). Single cell eQTL analysis identifies cell type-specific genetic control of gene expression in fibroblasts and reprogrammed induced pluripotent stem cells. bioRxiv, 2020.2006.2021.163766.

Okba, N., Müller, M., Li, W., Wang, C., GeurtsvanKessel, C., Corman, V., Lamers, M., Sikkema, R., de Bruin, E., Fd, C., et al. (2020). Severe acute respiratory syndrome coronavirus 2-specific antibody responses in coronavirus disease 2019 patients. Emerg Infect Dis.

Ou, X., Liu, Y., Lei, X., Li, P., Mi, D., Ren, L., Guo, L., Guo, R., Chen, T., Hu, J., et al. (2020). Characterization of spike glycoprotein of SARS-CoV-2 on virus entry and its immune cross-reactivity with SARS-CoV. Nature Communications 11, 1620.

Papatheodorou, I., Moreno, P., Manning, J., Fuentes, A.M., George, N., Fexova, S., Fonseca, N.A., Fullgrabe, A., Green, M., Huang, N., et al. (2020). Expression Atlas update: from tissues to single cells. Nucleic Acids Res 48, D77–D83.

Pavlovic, B.J., Blake, L.E., Roux, J., Chavarria, C., and Gilad, Y. (2018). A Comparative Assessment of Human and Chimpanzee iPSC-derived Cardiomyocytes with Primary Heart Tissues. Scientific reports 8, 15312.

Payne, D.C., Currier, R.L., Staat, M.A., Sahni, L.C., Selvarangan, R., Halasa, N.B., Englund, J.A., Weinberg, G.A., Boom, J.A., Szilagyi, P.G., et al. (2015). Epidemiologic Association Between FUT2 Secretor Status and Severe Rotavirus Gastroenteritis in Children in the United States. JAMA Pediatr 169, 1040–1045.

Pilotto, A., Odolini, S., Masciocchi, S., Comelli, A., Volonghi, I., Gazzina, S., Nocivelli, S., Pezzini, A., Foca’, E., Caruso, A., et al. (2020). Steroid-responsive severe encephalopathy in SARS-CoV-2 infection. medRxiv, 2020.2004.2012.20062646.

Qin, C., Zhou, L., Hu, Z., Zhang, S., Yang, S., Tao, Y., Xie, C., Ma, K., Shang, K., Wang, W., et al. (2020). Dysregulation of Immune Response in Patients With Coronavirus 2019 (COVID-19) in Wuhan, China. Clin Infect Dis 71, 762–768.

Ramani, A., Müller, L., Ostermann, P.N., Gabriel, E., Abida-Islam, P., Müller-Schiffmann, A., Mariappan, A., Goureau, O., Gruell, H., Walker, A., et al. (2020). SARS-CoV-2 targets cortical neurons of 3D human brain organoids and shows neurodegeneration-like effects. bioRxiv, 2020.2005.2020.106575.

Ramlall, V., Thangaraj, P., Meydan, C., Foox, J., Butler, D., May, B., de Freitas, J., Glicksberg, B.S., Mason, C., Tatonetti, N.P., et al. (2020). Identification of Immune complement function as a determinant of adverse SARS-CoV-2 infection outcome. medRxiv, 2020.2005.2005.20092452.

Rehbach, K., Zhang, H., Das, D., Abdollahi, S., Prorok, T., Ghosh, S., Weintraub, S., Genovese, G., Powell, S., Lund, A., et al. (2020). Publicly available hiPSC lines with extreme polygenic risk scores for modeling schizophrenia. bioRxiv, 2020.2007.2004.185348.

Ritchie, M.E., Phipson, B., Wu, D., Hu, Y., Law, C.W., Shi, W., and Smyth, G.K. (2015). limma powers differential expression analyses for RNA-sequencing and microarray studies. Nucleic Acids Res 43, e47.

Robinson, M.D., McCarthy, D.J., and Smyth, G.K. (2010). edgeR: a Bioconductor package for differential expression analysis of digital gene expression data. Bioinformatics 26, 139–140.

Samson, M., Libert, F., Doranz, B.J., Rucker, J., Liesnard, C., Farber, C.M., Saragosti, S., Lapoumeroulie, C., Cognaux, J., Forceille, C., et al. (1996). Resistance to HIV-1 infection in caucasian individuals bearing mutant alleles of the CCR-5 chemokine receptor gene. Nature 382, 722–725.

Schrode, N., Ho, S.M., Yamamuro, K., Dobbyn, A., Huckins, L., Matos, M.R., Cheng, E., Deans, P.J.M., Flaherty, E., Barretto, N., et al. (2019). Synergistic effects of common schizophrenia risk variants. Nat Genet 51, 1475–1485.

Shelton, J.F., Shastri, A.J., Ye, C., Weldon, C.H., Filshtein-Somnez, T., Coker, D., Symons, A., Esparza-Gordillo, J., Aslibekyan, S., and Auton, A. (2020). Trans-ethnic analysis reveals genetic and non-genetic associations with COVID-19 susceptibility and severity. medRxiv, 2020.2009.2004.20188318.

Shilts, J., and Wright, G.J. (2020). No evidence for basigin/CD147 as a direct SARS-CoV-2 spike binding receptor. bioRxiv, 2020.2007.2025.221036.

Si, L., Bai, H., Rodas, M., Cao, W., Oh, C.Y., Jiang, A., Moller, R., Hoagland, D., Oishi, K., Horiuchi, S., et al. (2020). Human organ chip-enabled pipeline to rapidly repurpose therapeutics during viral pandemics. bioRxiv, 2020.2004.2013.039917.

Song, E., Zhang, C., Israelow, B., Lu-Culligan, A., Sprado, A., Skriabine, S., Lu, P., Weizman, O.-E., Liu, F., Dai, Y., et al. (2020). Neuroinvasion of SARS-CoV-2 in human and mouse brain. bioRxiv, 2020.2006.2025.169946.

Su, S., Law, C.W., Ah-Cann, C., Asselin-Labat, M.L., Blewitt, M.E., and Ritchie, M.E. (2017). Glimma: interactive graphics for gene expression analysis. Bioinformatics 33, 2050–2052.

Takayama, K., Akita, N., Mimura, N., Akahira, R., Taniguchi, Y., Ikeda, M., Sakurai, F., Ohara, O., Morio, T., Sekiguchi, K., et al. (2017). Generation of safe and therapeutically effective human induced pluripotent stem cell-derived hepatocyte-like cells for regenerative medicine. Hepatol Commun 1, 1058–1069.

Tanigawa, Y., and Rivas, M. (2020). Initial Review and Analysis of COVID-19 Host Genetics and Associated Phenotypes. Preprints.

Thoms, M., Buschauer, R., Ameismeier, M., Koepke, L., Denk, T., Hirschenberger, M., Kratzat, H., Hayn, M., Mackens-Kiani, T., Cheng, J., et al. (2020). Structural basis for translational shutdown and immune evasion by the Nsp1 protein of SARS-CoV-2. Science 369, 1249–1255.

Toscano, G., Palmerini, F., Ravaglia, S., Ruiz, L., Invernizzi, P., Cuzzoni, M.G., Franciotta, D., Baldanti, F., Daturi, R., Postorino, P., et al. (2020). Guillain-Barre Syndrome Associated with SARS-CoV-2. N Engl J Med.

Tryka, K.A., Hao, L., Sturcke, A., Jin, Y., Wang, Z.Y., Ziyabari, L., Lee, M., Popova, N., Sharopova, N., Kimura, M., et al. (2014). NCBI’s Database of Genotypes and Phenotypes: dbGaP. Nucleic Acids Res 42, D975–979.

van der Made, C.I., Simons, A., Schuurs-Hoeijmakers, J., van den Heuvel, G., Mantere, T., Kersten, S., van Deuren, R.C., Steehouwer, M., van Reijmersdal, S.V., Jaeger, M., et al. (2020). Presence of Genetic Variants Among Young Men With Severe COVID-19. JAMA.

Walls, A.C., Park, Y.J., Tortorici, M.A., Wall, A., McGuire, A.T., and Veesler, D. (2020). Structure, Function, and Antigenicity of the SARS-CoV-2 Spike Glycoprotein. Cell.

Wang, K., Chen, W., Zhou, Y.-S., Lian, J.-Q., Zhang, Z., Du, P., Gong, L., Zhang, Y., Cui, H.-Y., Geng, J.-J., et al. (2020). SARS-CoV-2 invades host cells via a novel route: CD147-spike protein. bioRxiv, 2020.2003.2014.988345.

Wölfel, R., Corman, V.M., Guggemos, W., Seilmaier, M., Zange, S., Müller, M.A., Niemeyer, D., Jones, T.C., Vollmar, P., Rothe, C., et al. (2020). Virological assessment of hospitalized patients with COVID-2019. Nature.

Wrapp, D., Wang, N., Corbett, K.S., Goldsmith, J.A., Hsieh, C.L., Abiona, O., Graham, B.S., and McLellan, J.S. (2020). Cryo-EM structure of the 2019-nCoV spike in the prefusion conformation. Science 367, 1260–1263.

Wruck, W., and Adjaye, J. (2020). Meta-analysis of transcriptomes of SARS-Cov2 infected human lung epithelial cells identifies transmembrane serine proteases co-expressed with ACE2 and biological processes related to viral entry, immunity, inflammation and cellular stress. bioRxiv, 2020.2005.2012.091314.

Yan, R., Zhang, Y., Li, Y., Xia, L., Guo, Y., and Zhou, Q. (2020). Structural basis for the recognition of SARS-CoV-2 by full-length human ACE2. Science 367, 1444–1448.

Yang, L., Han, Y., Nilsson-Payant, B.E., Gupta, V., Wang, P., Duan, X., Tang, X., Zhu, J., Zhao, Z., Jaffre, F., et al. (2020). A Human Pluripotent Stem Cell-based Platform to Study SARS-CoV-2 Tropism and Model Virus Infection in Human Cells and Organoids. Cell stem cell.

Yang, N., Chanda, S., Marro, S., Ng, Y.H., Janas, J.A., Haag, D., Ang, C.E., Tang, Y., Flores, Q., Mall, M., et al. (2017). Generation of pure GABAergic neurons by transcription factor programming. Nat Methods.

Yuan, M., Wu, N.C., Zhu, X., Lee, C.-C.D., So, R.T.Y., Lv, H., Mok, C.K.P., and Wilson, I.A. (2020). A highly conserved cryptic epitope in the receptor-binding domains of SARS-CoV-2 and SARS-CoV. Science, eabb7269.

Zhang, R.R., Koido, M., Tadokoro, T., Ouchi, R., Matsuno, T., Ueno, Y., Sekine, K., Takebe, T., and Taniguchi, H. (2018). Human iPSC-Derived Posterior Gut Progenitors Are Expandable and Capable of Forming Gut and Liver Organoids. Stem Cell Reports 10, 780–793.

Zhang, T., Rodricks, M.B., and Hirsh, E. (2020). COVID-19-Associated Acute Disseminated Encephalomyelitis: A Case Report. medRxiv, 2020.2004.2016.20068148.

Zhang, Y., Pak, C., Han, Y., Ahlenius, H., Zhang, Z., Chanda, S., Marro, S., Patzke, C., Acuna, C., Covy, J., et al. (2013). Rapid single-step induction of functional neurons from human pluripotent stem cells. Neuron 78, 785–798.

Zhao, H., Shen, D., Zhou, H., Liu, J., and Chen, S. (2020a). Guillain-Barre syndrome associated with SARS-CoV-2 infection: causality or coincidence? The Lancet Neurology.

Zhao, J., Yang, Y., Huang, H., Li, D., Gu, D., Lu, X., Zhang, Z., Liu, L., Liu, T., Liu, Y., et al. (2020b). Relationship between the ABO Blood Group and the COVID-19 Susceptibility. medRxiv, 2020.2003.2011.20031096.

Zhou, J., Li, C., Liu, X., Chiu, M.C., Zhao, X., Wang, D., Wei, Y., Lee, A., Zhang, A.J., Chu, H., et al. (2020). Infection of bat and human intestinal organoids by SARS-CoV-2. Nat Med.

Zietz, M., and Tatonetti, N.P. (2020). Testing the association between blood type and COVID-19 infection, intubation, and death. medRxiv, 2020.2004.2008.20058073.

